# Visualization and Modeling of Inhibition of IL-1β and TNF-α mRNA Transcription at the Single-Cell Level

**DOI:** 10.1101/2020.10.16.342576

**Authors:** Daniel Kalb, Huy D. Vo, Samantha Adikari, Elizabeth Hong-Geller, Brian Munsky, James Werner

**Author notes:** These authors contributed equally to this work (Daniel Kalb, Huy D. Vo).

## Abstract

IL-1β and TNF-α are canonical immune response mediators that play key regulatory roles in a wide range of inflammatory responses to both chronic and acute conditions. Here we employ an automated microscopy platform for the analysis of messenger RNA (mRNA) expression of IL-1β and TNF-α at the single-cell level. The amount of IL-1β and TNF-α mRNA expressed in a human monocytic leukemia cell line (THP-1) is visualized and counted using single-molecule fluorescent in-situ hybridization (smFISH) following exposure of the cells to lipopolysaccharide (LPS), an outer-membrane component of Gram-negative bacteria. We show that the small molecule inhibitors MG132 (a 26S proteasome inhibitor used to block NF-κB signaling) and U0126 (a MAPK Kinase inhibitor used to block CCAAT-enhancer-binding proteins C/EBP) successfully block IL-1β and TNF-α mRNA expression. Based upon this single-cell mRNA expression data, we screened 36 different mathematical models of gene expression, and found two similar models that capture the effects by which the drugs U0126 and MG132 affect the rates at which the genes transition into highly activated states. When their parameters were informed by the action of each drug independently, both models were able to predict the effects of the combined drug treatment. From our data and models, we postulate that IL-1β is activated by both NF-κB and C/EBP, while TNF-α is predominantly activated by NF-κB. Our combined single-cell experimental modeling efforts shows the interconnection between these two genes and demonstrates how the single-cell responses, including the distribution shapes, mean expression, and kinetics of gene expression, change with inhibition.

## Introduction

Inflammation is a complex biological process that enables the host immune system to counteract potential biothreats. In the inflammatory response, select host receptors react to detrimental stimuli (e.g., pathogens, allergens, toxins, or damaged host cells), which activate various intracellular signaling pathways to secrete cytokines that trigger active recruitment of immune cells to the site of insult/infection.[1] While inflammation is usually beneficial to the host organism when fighting an infection, there is also a wide range of both chronic and acute conditions where remediation of inflammation is necessary for host recovery. For example, in certain viral infections, over-expression of inflammatory cytokines throughout the course of disease progression can lead to a potentially fatal cytokine storm that may be more harmful to the host than the underlying infection.[2] In addition to acute conditions, chronic inflammatory conditions, including rheumatoid arthritis, diabetes, [3] or persistent pain, can be caused by high concentrations of pro-inflammatory cytokines, such as Interleukin 1β (IL-1β) and Tumor Necrosis Factor α (TNF-α).

There are several drugs and medications used to limit or dampen the inflammatory response. The best known of these, non-steroidal anti-inflammatory drugs (NSAIDs), work by inhibiting the activity of cyclooxygenase enzymes (COX-1 and COX-2), which are important for the synthesis of key biological mediators and blood clotting agents.[4] Other drugs may act to inhibit key proteins involved in immune response signaling, such as kinase inhibitors or proteasome inhibitors. For kinase and proteasome inhibitors, these compounds are generally discovered first through binding assays, then studied in vitro by activity assays.[5-8] Cellular assays that monitor the effects of drugs in a more complicated environment generally follow such *in vitro* studies.[9] In a cellular assay, the effects of a drug can be studied by monitoring the level of inhibition of the target of interest, or may be studied by monitoring changes in a downstream signaling pathway. The role drugs play in dampening mRNA expression can be measured by quantitative PCR of the mRNA[10], through DNA microarrays[11], or by RNA sequencing.[12] While informative, most of these methods explore the response of large, ensemble populations of cells.

In contrast to traditional measurements of gene expression collected as bulk averages from large numbers of individual cells, single-cell techniques have revealed surprisingly rich levels of heterogeneity of gene expression.[13-16] When coupled with appropriate models, these distributions of single-cell gene expression can reveal fundamental information on expression kinetics and gene regulatory mechanisms, which is otherwise lost in the bulk measurements. [17, 18] Methods to measure gene expression in single cells generally rely on either amplification or imaging techniques. There are tradeoffs between the two techniques. Amplification-based methods, such as sequencing and PCR, provide high gene depth (tens-to-hundreds of genes can be analyzed) but can be expensive, generally analyze a small number of individual cells, and obscures spatial information.[19-23] Imaging methods generally utilize fluorescent oligonucleotide probes complementary to the RNA sequences of interest and include techniques such as single-molecule fluorescence *in situ* hybridization (smFISH)[16, 24, 25] and multiplexed barcode labeling methods.[26-28] Though fewer genes can be analyzed at one time, smFISH is relatively low cost, yields single-molecule resolution without the need for nucleic acid amplification, can readily measure several hundreds to thousands of individual cells, and directly visualizes the spatial location of each RNA copy.

There have been several studies that exploit single-cell methods in conjunction with single-cell modeling to study host inflammatory responses. For example, fluorescence flow cytometry was used to study the population switching between effector and regulatory T cells and to develop a computational model describing this dynamic behavior.[29] Application of single-cell RNA sequencing methods led to discovery of bimodal expression patterns and splicing in mouse immune cells.[30] Another study integrated live cell imaging and mathematical modeling to understand the ‘analog’ NF-κβ response of cell populations under ‘digital’ single-cell signal activation.[31] Additionally, a model of JAK1-STAT3 signaling was constructed following cell treatment by a JAK inhibitor with validation by wide field fluorescence microscopy.[32] In order to visualize single-cell immune responses, our lab previously used smFISH to monitor the single-cell mRNA expression of two cytokines, IL-1β and TNF-α, in a human monocytic leukemia cell line, THP-1, in response to lipopolysaccharide (LPS), a primary component of cell walls in Gram negative bacteria.[33] This work found a broad cell-to-cell heterogeneity in immune cell response to LPS.

Here, we exploit single-cell imaging and modeling methods to visualize and understand the broad distribution of mRNA responses to LPS stimulation in THP-1 immune cells. Moreover, these models were used to describe the effects of specific inflammatory inhibitors on the host immune response. The drugs employed, MG132, a 26S proteasome inhibitor used to block NF-κB signaling[8], and U0126, a MAPK kinase inhibitor known to block CCAAT-enhancer-binding proteins C/EBP[34], were selected for their differing roles in dampening the inflammatory response mediated by two key inflammatory cytokines: IL-1β and TNF-α. Our results show that MG132 inhibits both IL-1β and TNF-α mRNA expression, while U0126 primarily inhibits IL-1β expression. Models derived for the action of each drug independently can also accurately predict the behavior of the drug effects when applied in tandem. These results and models support the current biological understanding that IL-1β expression is activated by both NF-κB and C/EBP signaling pathways while TNF-α is predominantly activated by NF-κB. Notably, we observe that models developed to describe the effect single drugs can accurately predict the effect of drug combinations, paving the way for predictive computational analyses of combination drug therapies.

## Methods

### Microscopy and Image Analysis

A fully automated microscopy and image analysis routine was used to count and measure single-mRNA molecules as previously described.[33] In brief, a conventional wide-field microscope (Olympus IX71), arc lamp (Olympus U-RFL-T), high NA objective (Olympus 1.49 NA, 100X), 2D stage (Thorlabs BSC102), Z sectioning piezo (Physik Instrumente, PI-721.20) and cMOS camera (Hamamatsu orca-flash 4.0) are used to image single-cell mRNA content. Following image acquisition, a custom MATLAB script is used to: 1) automatically find and segment each individual cell based upon the bright-field cell image, the nuclear stain (DAPI), and the smFISH channel, 2) filter and threshold using a Laplacian-of-Gaussian filter (LOG) to find the single-mRNA copies, 3) fit all of the mRNA ‘spots’ to a 2D Gaussian using a GPU-accelerated algorithm, and 4) assign and count all single mRNA copies within each cell. Single-cell distributions are characterized by both their shapes and their mean values.

### Cell Culture

Human monocytic leukemia cells (THP-1, ATCC) were cultured in a humidified incubator with 5% CO2 at 37°C in R10% medium: RPMI-1640 Medium (with glutamine, no phenol red, Gibco) supplemented with 10% fetal bovine serum (FBS, ATCC). Cells were passaged every 5-7 days and used for experiments from age 60-120 days.

### Slide Preparation

Chambered cover-glass slides (#1.0 borosilicate glass, 8 wells, Lab-Tek) were coated with a sterile bovine fibronectin solution (1μg/well, Sigma, diluted in PBS, Gibco) overnight at 4°C. 10^5^ THP-1 cells/well were seeded onto fibronectin-coated slides for differentiation with R10% medium containing 100nM PMA (phorbol 12-myristate 13-acetate, Sigma) for 48hrs at 37°C. After differentiation, cells were serum-starved in serum-free RPMI-1640 Medium (no FBS) for 2hrs at 37°C. Cells were pre-treated with inhibitors (MG132, or U0126, or both) for 1hr at 37°C (10μM each in serum-free RPMI-1640 medium, 200μL/well). Untreated wells were kept in serum-free RPMI-1640 medium for 1hr at 37°C. Cells were then stimulated with a cocktail of 500μg/mL lipopolysaccharide (LPS, isolated from *E*.*coli* O55:B5, Sigma) and 10μM inhibitors (MG132, or U0126, or both) in R10% medium (200μL/well) for 30min, 1hr, 2hrs, or 4hrs at 37°C. Cells were washed in PBS and fixed in paraformaldehyde (4% solution in PBS (v/v), Alfa Aesar) for 15min at room temperature. Unstimulated cells were washed and fixed at t=0hrs immediately after 1hr inhibitor pre-treatment. After fixation, cells were washed twice in PBS, then permeabilized in 70% ethanol in RNase-free distilled water (v/v) (ThermoFisher) for at least 1hr (or up to 24 hrs) at 4°C. Cells were then washed in RNA FISH Wash Buffer A (Stellaris) for 20min at room temperature before RNA smFISH staining for mRNA.

### smFISH Staining for mRNA

Cells were stained with custom-designed RNA FISH probes (Stellaris) for IL-1β and TNF-α mRNA. Probes were diluted to 100nM each in RNA FISH Hybridization Buffer (Stellaris) containing 10% formamide (v/v) (ThermoFisher), then incubated on the fixed and permeabilized cells using 100μL/well for 4hrs at 37°C. Staining conditions were made in duplicate on each slide. Following probe hybridization, cells were washed three times in RNA FISH Wash Buffer A for 30min each time at 37°C, stained for 20min at 37°C with 100ng/mL DAPI solution (Life Technologies) in RNA FISH Wash Buffer A for 20min at 37°C, and washed in RNA FISH Wash Buffer B (Stellaris) for 20min at room temperature. Cells were washed once in PBS and stored in 200μL/well SlowFade Gold Anti-Fade Mountant (Life Technologies) diluted 4x in PBS for up to 7 days at 4°C. Unless otherwise specified, all steps were performed at room temperature, incubations were performed using 250μL/well, and washes were performed using 500μL/well.

### Stochastic reaction networks for modeling gene expression dynamics

The time-varying distributions of mRNA copy numbers observed from smFISH experiments are modeled in the framework of the chemical master equation (CME).[35, 36] This analysis proposes a continuous-time Markov chain in which each discrete state corresponds to a vector of integers that represents the copy number for each chemical species. In particular, we propose and compare different gene activation mechanisms with either two or three gene states and different ways in which the signal affects the gene activation/deactivation rates (see SI for details).

The probabilistic rate of a reaction event is determined through the propensity functions. The time-dependent probability vector *p(t)* over all states is the solution of the system of linear differential equations 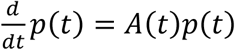, where *A(t)* is the transition rate matrix of the Markov chain. The CME was solved using the Finite State Projection (FSP) approach for marginal distributions.[37] All analysis codes are available at https://github.com/MunskyGroup/Kalb_Vo_2021.

### Conditionally independent models for simultaneous expression of multiple genes and in variable environmental conditions

The stochastic reaction network model above allows us to model the time-varying mRNA distribution for a *single* gene in a *single* experimental condition. However, our data comes with multiple genes and inhibitor treatment conditions, which necessitates a model to explain the joint mRNA count distribution of both IL-1β and TNF-α simultaneously. To do so, we make the assumption that the random variables describing IL-1β and TNF-α mRNA counts are *conditionally* independent, given a shared dependence on the same upstream time-varying NFκВ dynamics. These downstream gene expression variables are otherwise independent from each other, and can then be described by separated reaction networks that are coupled only by time, specific experiment condition, and the choice of parameters for the NFκВ reaction rates and the inhibitor effects. See SI for the precise mathematical description of this model.

### Parameter fitting

The full single-cell dataset consists of four independent biological replicates, each of which contains measurements of IL-1β and TNF-α mRNA copy numbers under four different inhibitor conditions (No inhibitor, with MG132, with U0126, and with both MG132 and U0126) and five measurement time points after LPS stimulation (0min (untreated), 30min, 1hr, 2hrs, and 4hrs). The ‘training’ dataset on which our CME model was parameterized consists of measurements made under three conditions (No inhibitor, with MG132, and with U0126). Parameters were estimated by minimizing the weighted sum of Kullback-Leibler divergences from the marginal empirical distributions of single-cell observations to those predicted by the CME model, which is equivalent to the log-likelihood of the observed joint distributions given the conditionally-independent model described in the previous section (See Section 3c in Supplementary Information). Evaluation of this likelihood requires the solutions of the CME, which were obtained using the Finite State Projection (FSP) algorithm.[37] (See SI for more details).

### Model evaluation and selection

We use a combination of statistical criteria to compare how well the different proposed mechanisms fit the ‘training’ data. These include the fit log-likelihood and the Bayesian Information Criteria (BIC, see Supplementary Information). In addition, we also compare the predictive performance of these alternative models using the log-likelihood of the dataset under the combined treatment that has both MG132 and U0126, which were not used for fitting the models.

## Results

### Inhibitor treatments reduce transcription levels of IL-1β and TNF-α in THP-1 human monocytic leukemia cells

The single-cell mRNA content of IL-1β and TNF-α in THP-1 cells were monitored over time after exposure to LPS and in response to two small molecule inhibitors MG132 and U0126, both alone and in combination. MG132 is a selective inhibitor of the NF-κB pathway, while U0126 inhibits the C/EBP pathway, part of the MAPK signaling cascade. [8, 38, 39] Representative images of gene expression after LPS exposure in the presence and absence of small molecule inhibitors are shown in Figures 1 and 2 for 1 and 2 hours post exposure.

**Figure 1.**
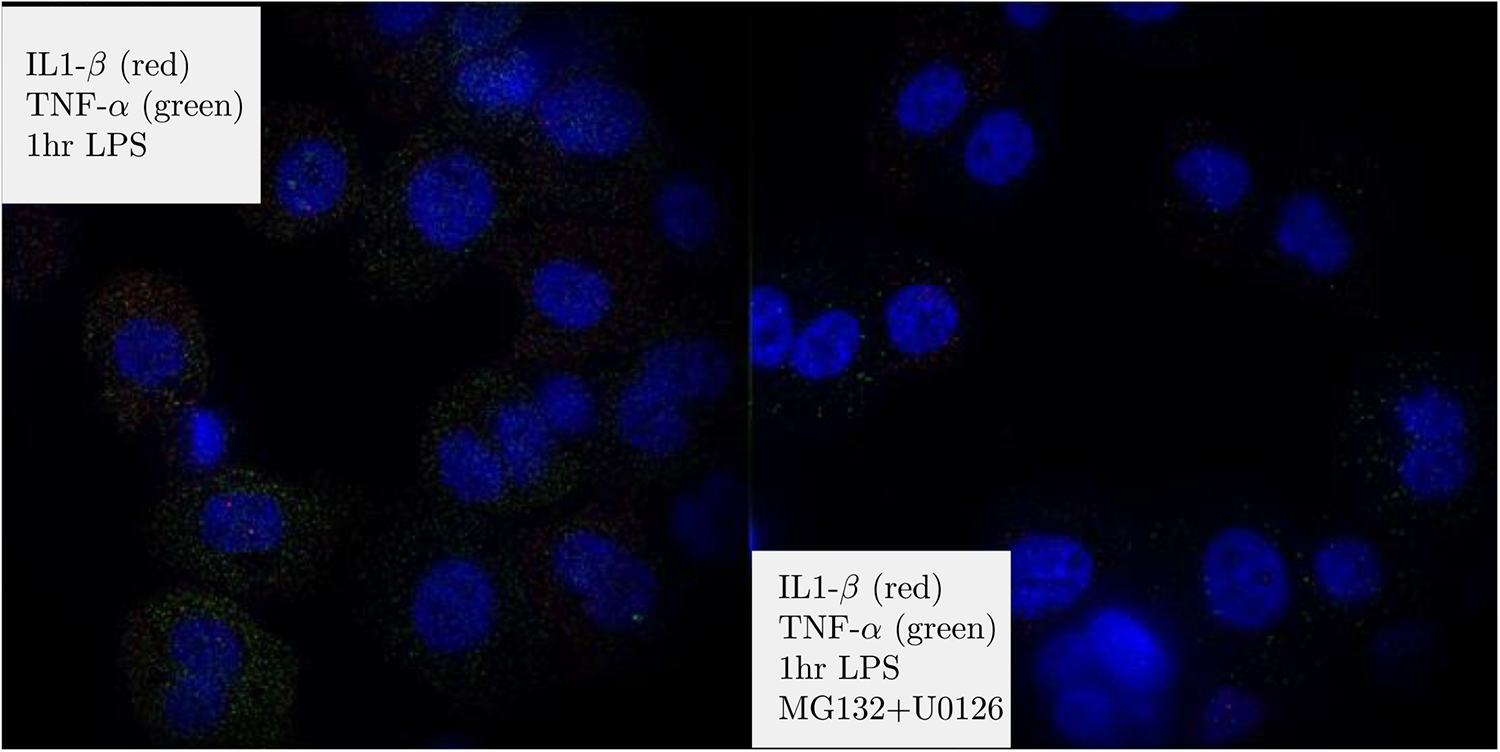
Representative images of IL-1β and TNF-α mRNA expression with and without inhibitors at 2 hrs LPS exposure. Images are LOG-filtered to emphasize single mRNA copies that appear as small diffraction limited spots in images. Blue is DAPI-stained nuclei, whereas red spots are individual copies of IL-1β and green spots are individual copies of TNF-α. Each image is ∼130 by 130 μm.

**Figure 2.**
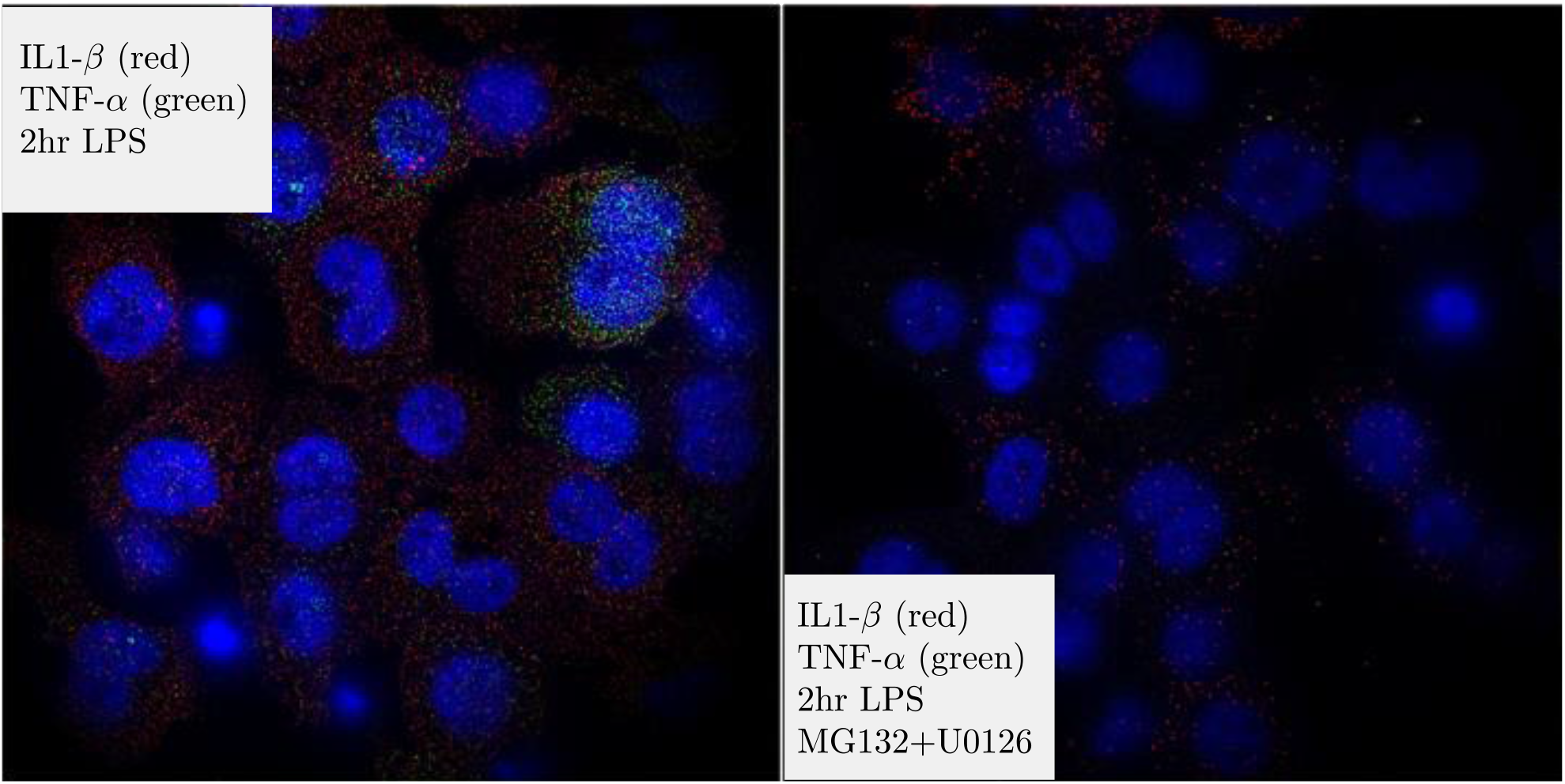
Representative images of IL-1β and TNF-α mRNA expression with and without inhibitors at 2 hrs LPS exposure. Images are LOG-filtered to emphasize single mRNA copies that appear as small diffraction limited spots in images. Blue is DAPI-stained nuclei, whereas red spots are individual copies of IL-1β and green spots are individual copies of TNF-α. Each image is ∼130 by 130 μm.

### IL-1β and TNF-α transcriptional responses to inhibitor conditions can be explained and predicted by signal-activated, multiple-state, stochastic bursting mechanisms

A class of several different two-state and three-state gene expression models were hypothesized to capture the stochastic transcriptional dynamics of the individual genes TNF-α and IL-1β (see Figure 3 for the schematics of these models). This class of model topologies has been used successfully in other works that examine MAPK-induced gene expression in single-cells.[40, 41] Here we present the interpretation of model ‘3SA’ in Figure 3 as an example (see Supplementary Information for the full list of reactions and parameters). In this model, each gene can exist in one of three transcriptional states: *G*_0_, *G*_1_, or *G*_2_. The biological interpretation of these states depends upon the specific parameter values chosen for the state’s transcription rate. For example, when the transcription rate in *G*_0_ is set to zero, then that could be thought of as an ‘off’ state; when the transcription rate in *G*_2_ is large, then that can be thought of as an ‘on’ state; and when the transcription in *G*_1_ takes an intermediate value, it could be described as a ‘ready’ or ‘poised’ state. The activation of each gene by C/EBP and NF-κB signals was modeled via time-dependent effects on gene-state transition rates. Specifically, for the model shown in Figure 3, the switching rate from *G*_0_ to *G*_1_ was assumed to depend on the time-varying abundance of NF-κВ, where NF-κВ increases the rate at which an ‘off’ gene switches to ‘ready’. More precisely, the time-dependent deactivation rate is given by *k*_01_*(t)* = *k*_01_ + *b*_01_[*NF − κB](t)*, where *k*_01_ is the basal gene activation rate and [*NF − κB](t)* is the concentration of NF-κВ, parametrized by a function of the form

**Figure 3.**
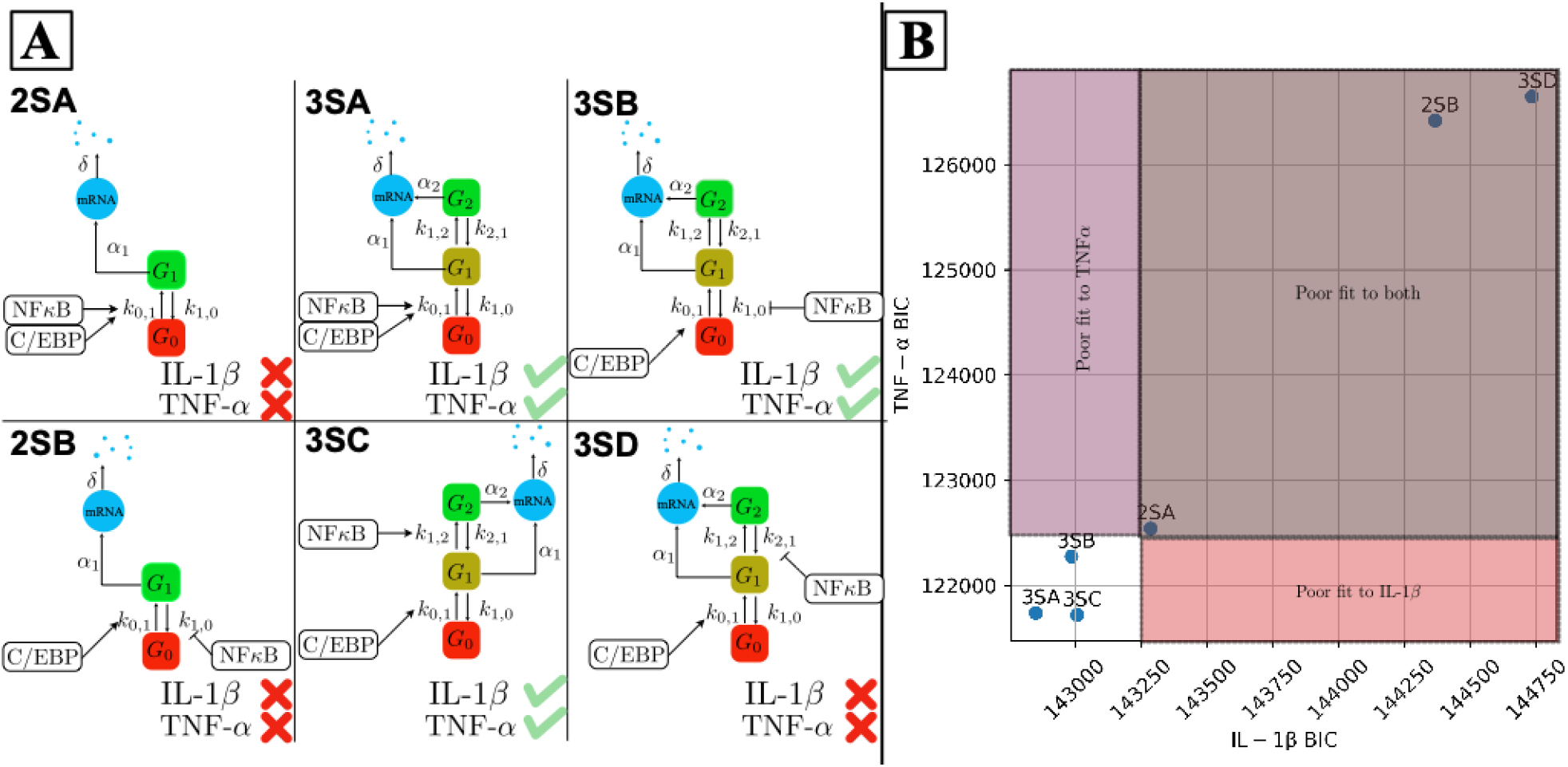
Signal-activated two- and three-state gene expression models considered for fitting the observed mRNA distributions. (A): Schematic diagrams of the six mechanisms considered. These models differ in the number of gene states and the mechanism by which NF-B increases the probability of transcription, either by increasing the rate of gene activation or inhibiting the rate of gene deactivation. (B): Performance evaluation of these models in terms of the Bayesian Information Criterion (BIC) based on IL-1β expression data and TNF-α expression data without inhibitor.

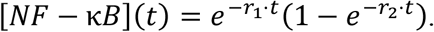

This model of NF-κВ activation is in general agreement with the literature on NF-κВ nuclear localization.[42] C/EBP was assumed to exert a constant influence on the rate of switching from *G*_0_ to *G*_1_. The expression dynamics of different genes (IL-1β, TNF-α) in response to different treatments (No inhibitors, with MG132, U0126, or both) were described by chemical master equations (CMEs) with the same reactions but different kinetic rate parameters. The effects of inhibitors, when present, were modeled as the reductions to the influence that C/EBP or NF-κВ exerted on gene activation.

We first attempt to independently fit the six gene expression models to the observed distributions of IL-1β and TNF-α, each individually under the inhibitor-free condition. Evaluating these fits using the Bayesian Information Criterion (BIC), we found that three of the different three-state models outperform all variants of the two-state models for both genes. From this comparison, we select these three variants of the three-state models and then extend them to postulate nine different model combinations (see Supplementary Figure 3), and we fit each of these models simultaneously to the mRNA distributions of both genes across all five time points and three experimental conditions (the remaining 27 combinatorial models that could have been constructed using one or more of the discarded models from above are ignored at this stage, although the effect of choice will be evaluated later). Specifically, we use the experimental data collected under inhibitor-free, MG132, and U0126 treatments to calibrate the parameters of all models. The full set of chemical reactions, as well as the fitted parameter values, are presented in the Supplementary Information. We then use the data under combined treatment (with both MG132 and U0126) as a testing dataset to see how well each of the fitted models predicts mRNA distributions under this experimental condition. The two models that yield the largest sum of the fit log-likelihood (computed on the training dataset under no or single inhibitor treatment) and the test log-likelihood (computed on the testing dataset under combined inhibitor treatment) are selected and illustrated in Figure 4. We confirm that these two models continue to provide good fits to the wildtype IL-1β and TNF-α expression data even when fitted simultaneously to both genes in all three conditions (Supplementary Figure 4). Moreover, since the final models continue to outperform the three previously discarded single-gene models for both genes in the drug-free condition, we are assured that these final models must also outperform all 27 of the discarded two-gene combinatorial models to match the drug-free data.

**Figure 4.**
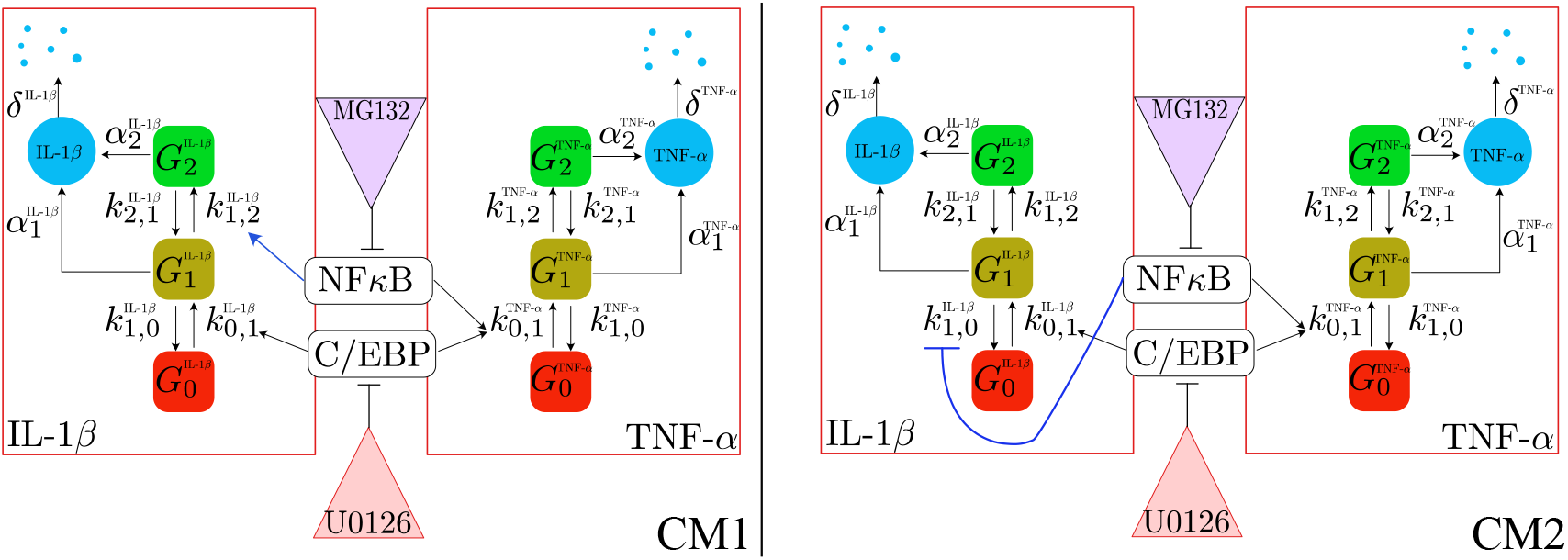
Two combinations of three-state gene expression models to simultaneously fit and predict the mRNA distributions transcribed from both IL-1β and TNF-α. These models are selected from a set of nine different combinations that can potentially explain the observed mRNA distributions in the experiment. In the first combined model (CM1), NF-κВ enhances the transition rate from G_1_ to G_2_ for the gene IL-1β. In the second model (CM2), NF-κВ inhibits the deactivation rate for IL-1β to switch from G_1_ to G_0_.

In the two combinatorial models selected (Figure 4), both mRNA species are transcribed via bursty mechanisms with three gene states. Both models suggest an identical gene expression mechanism for TNF-α, but provide different explanations for the expression of IL-1β. Specifically, they differ in how the effects of NF-κВ signal on IL-1β gene activation are explained. The first combinatorial model (CM1) postulates that the presence of NF-κВ enhances the transition rate for IL-1β from *G*_1_ to *G*_2_ by an additive term proportional to NF-κВ concentration in the nucleus. On the other hand, the second combinatorial model (CM2) postulates that the same signal inhibits the deactivation rate of IL-1β for transiting from *G*_1_ to *G*_0_. In either case, the activity of NF-κВ leads to a greater chance that the gene moves from the ‘off’ state and through the ‘ready’ state to reach the ‘on’ state.

Figure 5 shows the single-cell mRNA distribution shapes of both IL-1β and TNF-α in response to LPS as well as the best combined model fit to these data, and Supplementary Figures 6 and 7 show expanded results for the fits of both genes in all time points and conditions. For both genes, the data are indicative of ‘bursting’ gene expression, characterized by most cells exhibiting lower expression and a long ‘tail’ of relatively rare high-expressing cells. The distributions of mRNA copies per cell in the presence of the small-molecule inhibitors (Figure 5(B-D) and Figure 5(F-H)) retain their bursting shape (similar to the expression patterns seen with no LPS (Figure 5A and Figure 5E)). Based upon how the cell-to-cell mRNA distributions change in the presence of the drugs MG132 and U0126, we postulated that these drugs could modulate how NF-κВ or C/EBP regulate gene expression. We note that kinetic parameters were determined from the measured mRNA distributions in the drug free and single-drug exposure time-course experiments (Table 1). These parameters then were used to predict the combined drug condition (without any additional fitting of the data), yielding a good approximation of the measured mRNA distributions (Figure 5 D,H and bottom rows of Supplementary Figures 6 and 7).

**Figure 5.**
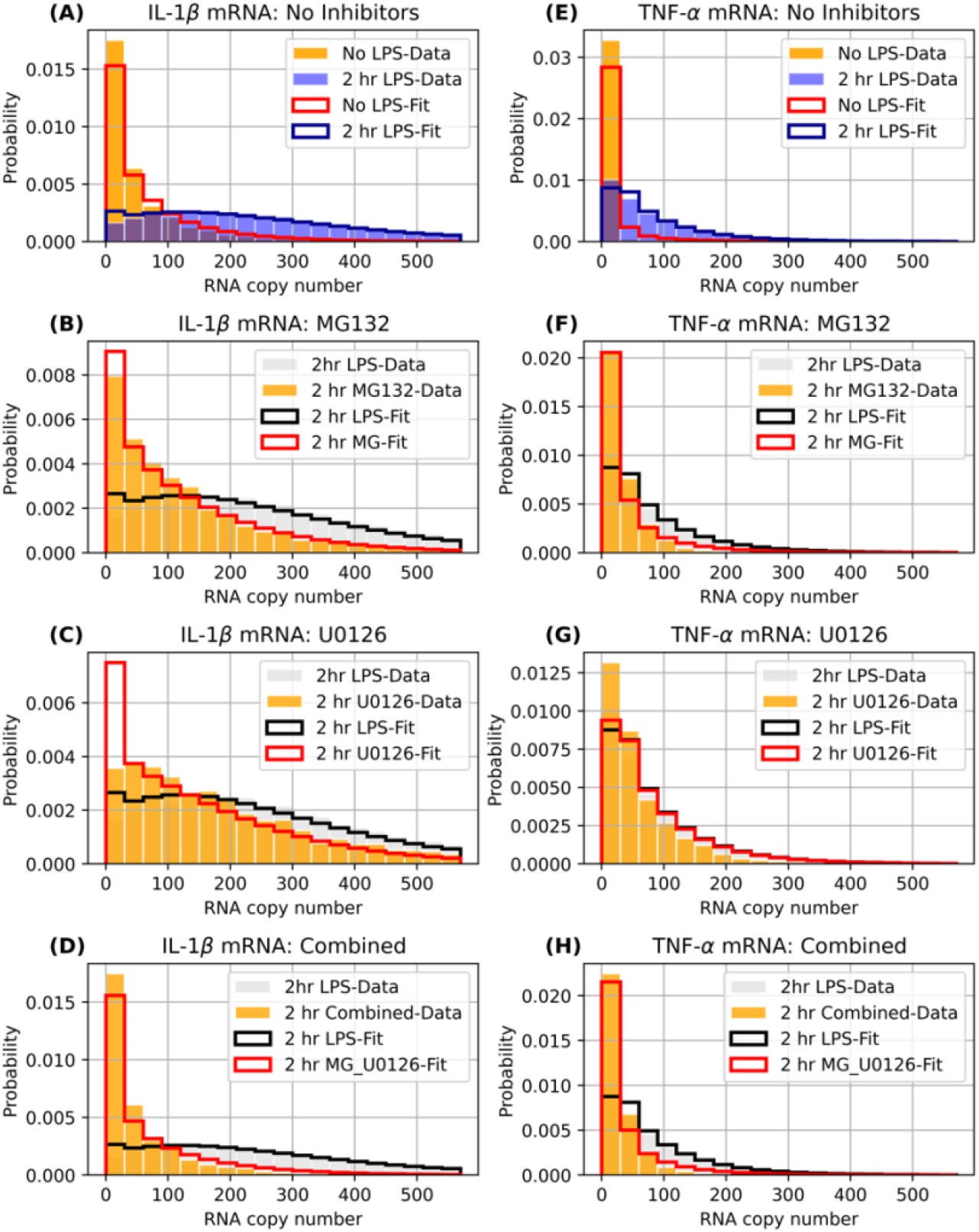
Distributions for single-cell mRNA content. (A-D) Probability distribution (data represented as bars, model fits/predictions as solid lines) for number of IL-1β copies per cell with: (A) 2hr LPS exposure with no inhibitor treatment, (B) 2hr LPS exposure with MG132, (C) 2hr LPS exposure with U0126, and (D) 2hr LPS exposure with U0126 and MG132 combined. (E-H) same as (A-D), but for TNF-α. For reference, each panel shows the corresponding mRNA distribution at 2hr LPS exposure with no inhibitor treatment (data in grey, model in black). A-C and E-H show the model fits to data with no inhibitors or a single inhibitor, and D,H show the validation of model predictions for the two inhibitor combination.

**Table 1.** Fitted parameters for the two best performing combined multi-gene, multi-condition gene expression models. These are obtained by fitting the model-predicted distributions of RNA copy number to data collected under three inhibitor conditions (no inhibitors, MG132, and U0126).

|  | CM1 |  | CM2 |  |  |
| --- | --- | --- | --- | --- | --- |
| Parameter | IL-1β | TNF-α | IL-1β | TNF-α | Interpretation |
| $r_1$ | 9.01e-04 | | 1.03e-03 | | Parameters for NF-kB dynamics (second<sup>-1</sup>) |
| $r_2$ | 3.05e-04 | | 4.39e-04 | | |
| $k_{01}$ | 3.89e-02 | 1.21e-04 | 4.77e-03 | 1.17e-04 | LPS-free transition rate $G_0$ to $G_1$ (second<sup>-1</sup>) |
| $b_{01}$ | NA | 2.27e-02 | NA | 2.09e-02 | Multiplicative factor for NF-kB induced increase in gene activation rate (second<sup>-1</sup>) |
| $k_{10}$ | 7.62e-03 | 4.67e-02 | 7.66e-02 | 5.48e-02 | LPS-free gene deactivation rate $G_1$ to $G_0$ (second<sup>-1</sup>) |
| $b_{10}$ | NA | NA | 6.60e+00 | NA | Multiplicative factor for NF-kB induced decrease in gene deactivation ($G_1$ to $G_0$) rate (second<sup>-1</sup>) |
| $k_{12}$ | 3.93e-05 | 8.19e-03 | 5.50e-04 | 9.41e-03 | LPS-free transition rate $G_1$ to $G_2$ (second<sup>-1</sup>) |
| $b_{12}$ | 9.60e-03 | NA | NA | NA | Multiplicative factor for NF-kB induced increase in transition rate $G_1$ to $G_2$ (second<sup>-1</sup>) |
| $k_{21}$ | 8.37e-03 | 3.98e-03 | 8.79e-03 | 4.03e-03 | Transition rate from highly activated state to moderately activated state ($G_2$ to $G_1$) (second<sup>-1</sup>) |
| $\alpha_0$ | 1.09e-04 | 3.61e-05 | 5.29e-05 | 2.60e-05 | Basal transcription rate when gene is at basal state $G_0$ (molecule/second) |
| $\alpha_1$ | 1.64e-05 | 6.24e-01 | 1.00e-06 | 7.07e-01 | Transcription rate when gene is at $G_1$ (molecule/second) |
| $\alpha_2$ | 9.99e-01 | 5.39e-01 | 1.00e+00 | 5.41e-01 | Transcription rate when gene is at $G_2$ (molecule/second) |
| $\delta$ | 5.67e-05 | 2.29e-04 | 5.27e-05 | 2.16e-04 | mRNA degradation rate (molecule/second) |
| $b_{01}^{MG}$ | NA | 7.80e-03 | NA | 7.18e-03 | MG-modulated value of $b_{01}$ (second<sup>-1</sup>) |
| $b_{10}^{MG}$ | NA | NA | 6.80e-01 | NA | MG-modulated value of $b_{10}$ (second<sup>-1</sup>) |
| $b_{12}^{MG}$ | 2.83e-03 | NA | NA | NA | MG-modulated value of $b_{12}$ (second<sup>-1</sup>) |
| $k_{01}^{U0126}$ | 2.22e-03 | 1.00e-04 | 2.22e-03 | 1.00e-04 | U0126-modulated value of $k_{01}$ (second<sup>-1</sup>) |

### Measurement and analysis of mRNA expression suggests that MG132 inhibits both TNF-α and IL-1β, while U0126 inhibits only IL-1β

Figure 6 shows the measured and model-predicted mean (Figures 6A-B) and standard deviation (Figures 6 C-D) for mRNA expression versus time from 0 to 240 min post LPS exposure for both genes. In the absence of inhibitors, the cells show a rapid increase in both IL-1β and TNF-α mRNA content following introduction of LPS, with expression peaking at ∼325 IL-1β mRNA copies per cell at 2 hrs and at ∼200 TNF-α mRNA copies per cell at 1 hr. In the presence of inhibitors, we see that IL-1β expression is inhibited by both U0126 and MG132, but with different kinetic trajectories. MG132 treatment dampens IL-1β expression with maximum expression at ∼140 mRNA copies per cell at 1 hr. In contrast, U0126 strongly inhibits IL-1β expression at early time points (0-30 min), but is less effective at later time points, with maximal expression at ∼200 mRNA copies per cell after 120 min. The combination of the two inhibitors results in low expression of IL-1β across all time points, with <50 mRNA copies per cell. For TNF-α, MG132 markedly reduces expression from ∼200 mRNA copies per cell to <50 mRNA copies per cell at 60 min. Interestingly, U0126 shows very little inhibition of TNF-α when used alone. Addition of both inhibitors led to low expression of TNF-α, similar to MG132 treatment alone. In the rest of this section, we will provide a more detailed explanation of these observations based on the model fits.

**Figure 6.**
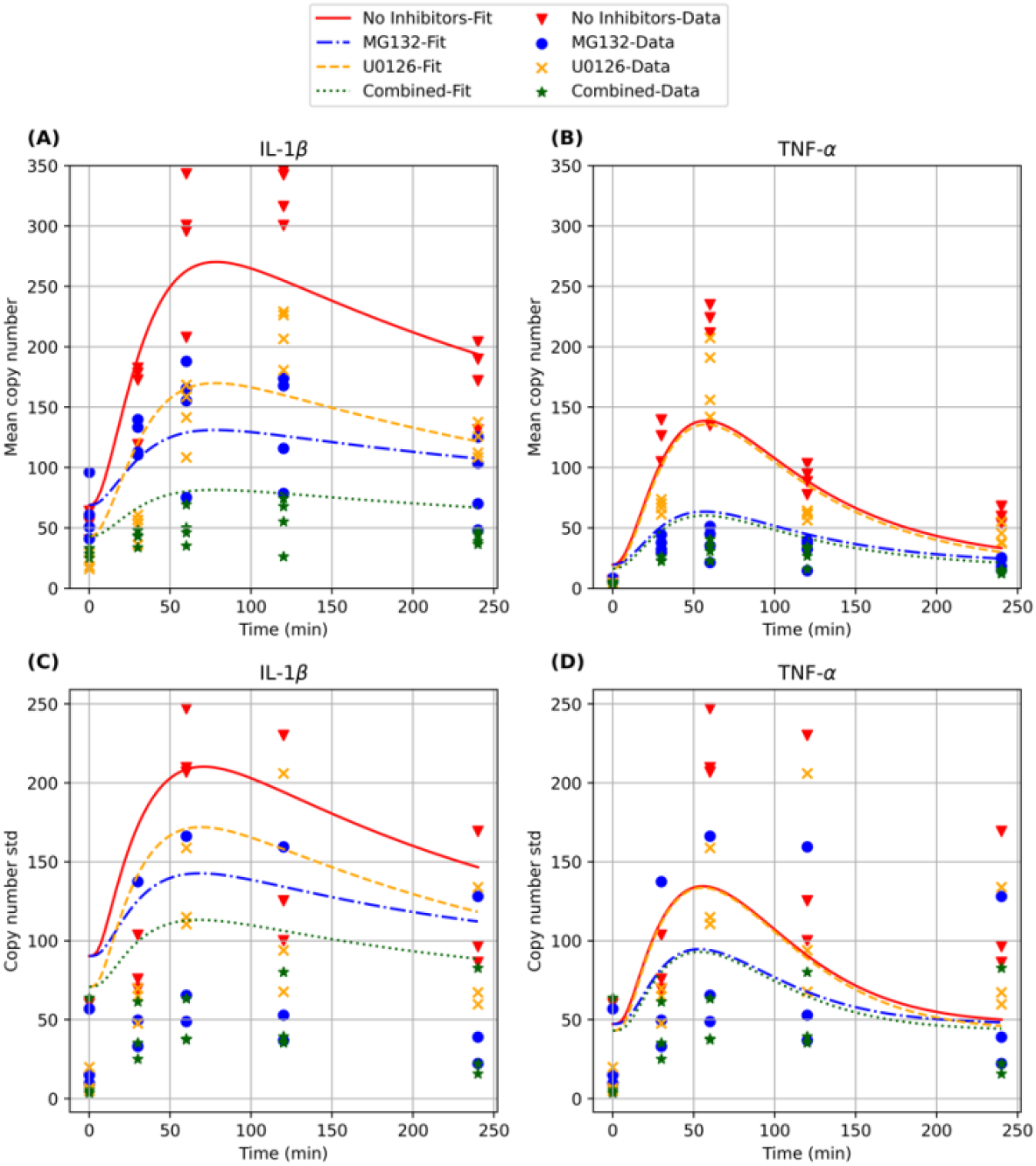
Mean and standard deviation of mRNA copy numbers per cell with and without inhibitors over 4hrs of LPS exposure (timepoints 0min, 30min, 1hr, 2hr, and 4hr) estimated from four independent biological replicates per inhibitor condition (markers) and model fits based on the combinatorial model CM1 (solid and dashed lines). **(A)&(B**): mean mRNA copy numbers per cell for IL-1β and TNF-α. **(C)&(D):** standard deviations of mRNA copy numbers per cell for IL-1β and TNF-α. See Supporting Information (section 1b) for details on our computation of these model-predicted statistics.

### The selected three-state gene expression models provide descriptive explanations for the activation dynamics of IL-1β and TNF-α under LPS stimulation

The two best-fit models allow us to propose several mechanisms for signal-activated expression of IL-1β and TNF-α, as well as how these mechanisms are affected by the small-molecule inhibitors MG132 and U0126. The inferred dynamics of NF-κВ concentration, which are not directly observed from data, is qualitatively similar between two models (Figure 7A). For TNF-α, both models yield similar fitted parameters that lead to identical interpretation. In the absence of LPS, the deactivation rate *k*_10_ for TNF-α is about 385 times higher than the activation rate *k*_01_ for model CM1 (similar comparison for CM2). As a consequence, the gene spends most of its time in the basal state that has a very low basal mRNA production rate (∼ 10^−5^ molecules per second). Under LPS stimulation, NF-κВ concentration in the nucleus quickly increases to its maximal value in about 15 minutes, with the downstream effects of increasing the fractions of cells in the active states (Figure 7C and 7E). As a consequence, there is a temporary increase in the mean mRNA production rate (Figure 7G), which explains the increased width of the distribution of TNF-α mRNA copy numbers observed. The signal starts decaying shortly after reaching its peak at around 15 minutes, resulting in less mRNA being produced and the mean TNF-α copy number slowly decreases (Figure 6B).

**Figure 7.**
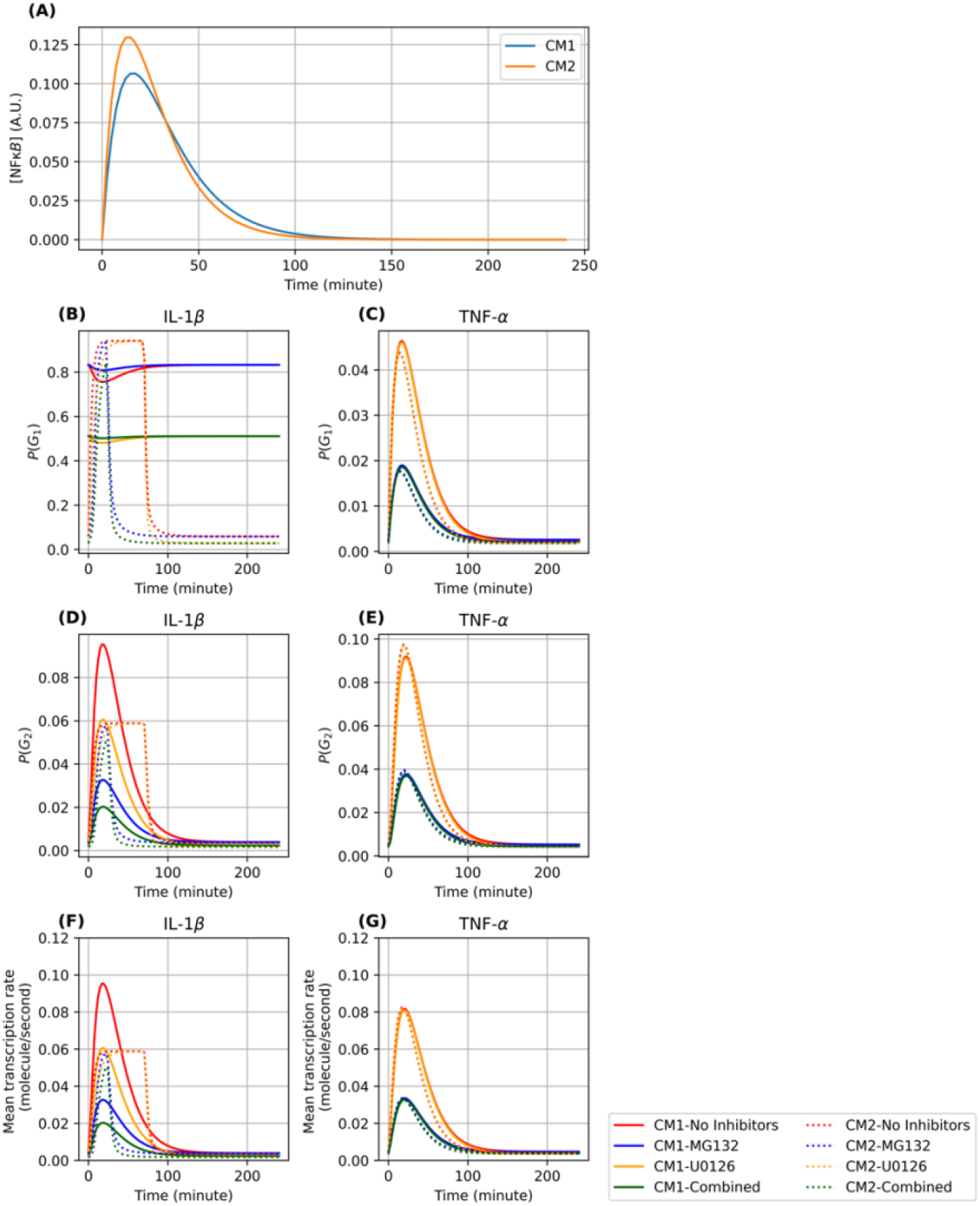
Model-predicted downstream influence of NF-κВ using the best two models. **(A):** Signal strength of NFκВ in the nucleus in arbitrary units (AU). **(B-C):** The time-varying probability of IL-1β and TNF-α to occupy the intermediate gene state (G_1_). **(D-E):** The time-varying probability of IL-1β and TNF-α to occupy the final gene state (G_2_). **(F-G):** The time-varying mean transcription rates for IL-1β and TNF-α, under four different inhibitor conditions (No inhibitors, MG132, U0126, both MG132 and U0126).

For IL-1β, both models produce similar mRNA copy number distributions at the time points where experimental measurements were taken across experimental conditions (Supplementary Figure 6). In addition, parameter fits for both models suggest that mRNA transcription rates are low when IL-1β is in the states *G*_0_ and *G*_1_ (of the order of 10^−4^ and 10^−5^ respectively in model CM1, and 5 × 10^−5^ and 10^−6^ respectively in model CM2), while the transcription rate at state *G*_2_ is high (both models fit to approximately one molecule per second). These fits allow us to interpret for IL-1β the state *G*_1_ as an intermediate “permissive” state from which the gene can become fully active at *G*_2_. Prior to LPS stimulation, the rate at which IL-1β switches to the fully active state *G*_2_ from the intermediate state *G*_1_ is about 212 times smaller than the reverse rate in model CM1 (and 15 times smaller in model CM2). As a consequence, IL-1β stays in the basal state for most cells, with model CM1 suggesting that a significant fraction of the cells are in the intermediate state without switching over to the highly active state *G*_2_, whereas model CM2 sugests that the fractions of cells in *G*_1_ and *G*_2_ are both low (Figure 7B,D). As a consequence, IL-1β stays in the basal states for most cells. Upon LPS induction, however, the increased NF-κВ concentration either has a positive effect on the rate for switching from *G*_1_ to *G*_2_ (model CM1), or an inhibitory effect on the deactivation rate from *G*_1_ to *G*_0_. This either allows for more cells already in the intermediate state *G*_1_ to switch to the active state *G*_2_ (model CM1), or for a significant increase in the fraction of cells in state *G*_1_ that consequently switch to *G*_2_ (model CM2) (Figure 7B). Either way, IL-1β has markedly higher probability to be in the fully activated state *G*_2_ (Figure 7D), leading to an increase in the mean IL-1β mRNA transcription rate. This increased production is sustained for a relatively short time but achieves a high maximal value in model CM1, while it is sustained for longer but with a lower maximum in CM2. Despite these differences for the intermediate, unobserved, components, both models yield fits for IL-1β degradation rates whose relative differences are below ten percent (5.67 × 10^−5^ molecules/second in CM1 and 5.27 × 10^−5^ molecules/second in CM2). The longer half-life of IL-1β compared to TNF-α also explains why mean IL-1β mRNA levels remain higher than TNF-α despite both genes reverting to their basal levels as NF-κВ fades away at about 100 minutes.

### The selected models suggest that NF-κВ activates both TNF-α and IL-1β, while C/EBP has no major influence on TNF-α transcriptional activity

In addition to providing explanations for IL-1β and TNF-α transcriptional dynamics under LPS, our exhaustive search for the reaction network parameters (Table 1) also leads to a quantitative understanding of the effects of the small-molecule inhibitors MG132 and U0126 on these genes. In the presence of the inhibitor MG132, the activation effect of NF-κВ is substantially reduced for both genes (for model CM1, the ratio 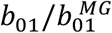 is about 2.9 for TNF-α and 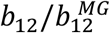 is about 3.4 for IL-1β; for model CM2, the ratio 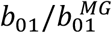 is about 2.9 for TNF-α and 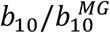 is about 9.7 for IL-1β), leading to smaller fractions of cells in the active states and consequently lower overall TNF-α mRNA production. The inhibitor U0126 decreases the activation rate *k*_01_ of IL-1β by 2.18 fold in CM1 and 4 fold in CM2, leading to overall reduction in IL-1β transcription. On the other hand, the addition of U0126 only reduces the rate *k*_01_ from 1.21 × 10^−4^ to 1.00 × 10^−4^ in model CM1 and from 1.17 × 10^−4^ to 1.00 × 10^−4^ in model CM2. Since U0126 is known to inhibit C/EBP, this suggests that C/EBP does not have a major influence on TNF-α.

## Discussion

Single-cell measurements allow for a more complete description and characterization of gene expression kinetics and regulatory mechanisms. Applying single-cell measurement techniques, we have demonstrated that gene transcription heterogeneity of two key immune response genes, IL-1β and TNF-α, occurs to a surprising extent within a seemingly uniform cell population following an immune assault. Such measurements of cell-to-cell distribution can be more informative than average values obtained from bulk measurements. For example, cells in the tail of the distribution may ultimately dictate the fate of disease progression rather than the average response, such as in the case of highly stimulated cells that lead to a cytokine storm. This is analogous to understanding how certain individuals (such as super-spreaders) may dictate the pathway of an epidemic more than basic reproduction number (R0) values.[43] Here we show that IL-1β and TNF-α genes, while upregulated upon bacterial LPS exposure, can be suppressed at the transcriptional level by the inhibitors MG132 and U0126. Interestingly, each of these inhibitors has a different kinetic effect. U0126 inhibits early IL-1β expression, while MG132 causes a delayed inhibition pattern, suggesting that C/EBP signaling occurs prior to NF-κB activity, in response to LPS (see Figure 6). Additionally, we show that TNF-α is predominantly and rapidly inhibited by MG132 treatment, suggesting that NF-κB is the primary upstream regulator of TNF-α expression in response to LPS.

To describe these results, we considered 36 potential stochastic models to reproduce IL-1β and TNF-α activity, and we found that the time-course of the cell-to-cell distributions of transcript copy numbers for both Il-1β and TNF-α could be adequately captured by two stochastic models, each having three states for gene transcription. Moreover, the effects of anti-inflammatory drugs MG132 and U0126 on the mRNA copy numbers of these two genes could be captured with both three state models. While the kinetic models were fit to data for each drug acting independently, both models were able to predict the data well for the combined drug treatment. The final two models selected identical mechanisms and dynamics for the regulation of TNF-α activity, but different mechanisms for the control IL-1β. Interestingly, although both models make indistinguishable predictions for the distributions of mature IL-1β mRNA in all conditions and time points measured for this study (Supplementary Figure S6 and S7), the two models make qualitatively and quantitatively different predictions for other, as yet untested experimental conditions. Specifically, the two models differ in their predictions for the instantaneous transcription rate at early times, where model CM1 predicts a short period of high transcription activity and model CM2 predicts a sustained period of moderate strength activity (Figure 7F). Furthermore, the two models also differ how the instantaneous transcription rate would be affected by MG132 treatment. In principle, our analyses suggest that these two models could be resolved using intron smFISH labeling to measure nascent transcription activity to quantify instantaneous transcription rates in shorter time scale experiments (e.g., 40 to 80 minutes). These experiments are beyond the scope of the current study and are left for future investigation.

Our results suggest that the integration of single-cell measurements and predictive kinetic modeling can lead to improved mechanistic understanding that could eventually lead to more effective combination therapies against chronic and acute inflammatory diseases. We note that while there are possible ways to extend the model proposed in this study to describe the joint expression of both IL-1β and TNF-α, the large state space required to analyze the joint expression of more than two mRNA species, coupled with the complexity of integrating time-varying kinase signals, poses a prohibitive challenge for current computational tools. Advances in high performance FSP-based inference methods (e.g., [44]) may potentially allow us to tackle the joint modeling approach in future work. Overall, this study emphasizes the need for further use of single-cell measurements to understand gene responses in order to identify outlier cells and capture full distributions.[45] Single-cell gene expression measurements combined with the appropriate model could provide otherwise overlooked insights into the kinetics, spatial distribution, and regulatory mechanisms of any number of genes.

## Acknowledgements

DK, SA, EH-G, and JHW were supported by the Los Alamos Directed Research and Development (LDRD) program. HV and BM were supported by National Institutes of Health under grant R35 GM124747. This work was performed, in part, at the Center for Integrated Nanotechnologies, an Office of Science User Facility operated for the U.S. Department of Energy (DOE) Office of Science. Los Alamos National Laboratory, an affirmative action equal opportunity employer, is managed by Triad National Security, LLC for the U.S. Department of Energy’s NNSA, under contract 89233218CNA000001.

