## Supplemental Information for "Visualization and Modeling of Inhibition of IL-1β and TNF-α mRNA Transcription at the Single-Cell Level"

Supplementary information for "Visualization and Modeling of IL1β and TNFα mRNA Transcription at the Single-Cell Level"

### 1) Stochastic reaction network and the chemical master equation

#### a) Markov modeling of stochastic biochemical reactions

We model the stochastic transcriptional dynamics of IL-1β and TNFα using a *stochastic reaction network* (SRN) [1], which is a class of Markov jump processes in which each state is a vector $\boldsymbol{x} = (x_1, \dots, x_N)^T$ of integral copy numbers, corresponding to the $N$ molecular species in the network, in which transitions represent the chemical reactions between these species. Reaction events are assumed to occur at discrete times, with independent waiting times between two consecutive reaction events. Upon the firing of reaction channel $R_j$ $(j = 1, \dots, M)$, the state $\boldsymbol{x}$ is perturbed by an amount determined by the stoichiometry vector $\nu_j$ associated with $R_j$. The relative likelihood for each reaction channel to fire is characterized by propensity functions $\alpha_j(t,x), j = 1, \dots, M$.

Since we want to model smFISH data, which consists of independent single-cell population snapshots measured at different time points, it is necessary to characterize the probability distribution of the state $\boldsymbol{x}$ at specific time points. In particular, let $p(t)$ denote the probability distribution of the states at time $t$, then $p(t)$ is the solution of the chemical master equation (CME)

$$\frac{d}{dt}p = A(t)p(t).$$

Here, $A(t) = \left[a_{x,y}(t)\right]_{x,y}$ is the time-dependent transition rate matrix of the Markov chain indexed by the states, with $a_{x,y}(t) = \alpha_j(t,y)$ if $x = y + \nu_j$, $a_{x,x}(t) = -\sum_{z\neq x} a_{z,x}(t)$, and $a_{x,y} = 0$ otherwise.

#### b) Finite State Projection algorithm for computing temporally-varying single-cell distributions and their first and second-order moments

We use an in-house C++ implementation [2] of the finite state projection (FSP) algorithm [3] to directly compute the solution of the CME. In essence, the FSP is based on approximating the infinite-state Markov process of the SRN with a finite-state Markov process, in which all states outside of a chosen finite subset $J$ are aggregated into a sink state $\boldsymbol{G}$. This effectively reduces the CME into a finite set of ordinary differential equations (ODEs) that can be solved by existing numerical solvers. The finite solution $p_J(t)$ is then used to approximate the true solution $p(t)$. The probability concentrated at the sink state $\boldsymbol{G}$ provides the exact quantification of the total variation distance between the approximate and true solutions, which we utilize to adaptively expand the state space to always keep the error below $10^{-4}$.

The outputs $p_J(t)$ of the FSP also provide us approximations for the mean and variance of mRNA copy numbers shown in Figure 6 in the main text. These are done straightforwardly by computing the first- and second-order moments of the FSP distribution $p_J(t)$.

### 2) Multiple-state gene expression models for IL-1β and TNFα transcription

#### a) Basal gene expression dynamics

The first step in our modeling approach is to specify a model for the transcription dynamics of each gene IL-1β and TNF-α in the absence of LPS-induced signals. We refer to this model as the *basal single-gene model*. The model keeps track of the state vector $\boldsymbol{x} = (G_0, \dots, G_n, R)$ where $G_i$ is one if the gene is in state $i$ and zero otherwise. The gene states are reachable from each other via a linear chain of reversible reactions $G_0 \rightleftharpoons \cdots \rightleftharpoons G_n$. Once the gene leaves the inactive state, it can be transcribed into mRNA molecules. The component $R$ in the state vector keeps track of the number of mRNA present in the system. The full set of reactions are:

$$G_0 \to G_1,$$

$$G_1 \to G_0,$$

$$\vdots$$

$$G_{n-1} \to G_n,$$

$$G_i \to G_i + R, i = 0, \dots, n,$$

$$R \to \emptyset.$$

b) **Gene expression dynamics under LPS stimulation without inhibitor**

In the next step, we modify the basal models to describe the stochastic dynamics of each gene in the presence of LPS induction, but without any inhibitors. We assume that the LPS-induced relative abundance level of NF-kB molecules takes the form

$$S_{\text{NF-κB}}(t) = \exp(-r_1 t)\,(1 - \exp(-r_2 t)). \tag{1}$$

where $t$ is the time after LPS stimulation. In addition, we assume that NF-kB modulates one and only one of the transition rates between gene states. The transition rate from $G_0$ to $G_1$, $k_{01}$, is proportional to C/EBP concentration, which is assumed to be constant for each inhibitor condition.

From the basic template just described, we then propose different mechanisms of gene expression dynamics as illustrated in Figure 3 in the main text. Specifically, depending on whether there are $n = 2$ or $n = 3$ gene states, and whether NF-kB concentration affects the rate of the gene state transition (A) $G_0 \to G_1$, (B) $G_1 \to G_0$, (C) $G_1 \to G_2$, or (D) $G_2 \to G_1$ (whenever such transition exists in the network).

Whenever NF-κB affects the transition $G_i \to G_{i+1}$ (as in cases A and C), the rate is assumed to be time-dependent and takes the form

$$k_{i,i+1}^{TV}(t) = k_{i,i+1} + b_{i,i+1} S_{\text{NF-κB}}(t),$$

where $i \in \{0, 1\}$ and $a_{i,i+1}, b_{i,i+1}$ are assumed to be positive. On the other hand, when NF-κB affects the reverse transitions $G_{i+1} \to G_i$ (as in cases B and D), the transition rates assume the form

$$k_{i+1,i}^{TV}(t) = \max\{\,0, k_{i+1,i} - b_{i+1,i} S_{\text{NF-κB}(t)}\}.$$

We refer to the parameters $b_{i,i+1}$ (in case A or C) or $b_{i+1,i}$ (in case B or D) as the *signal strength* parameters, since the larger the values of these parameters, the stronger NF-κB concentration affects gene state switching rates compared to the basal levels. To sum up, we propose six reaction network structures that we label as 2SA, 2SB, 3SA, 3SB, 3SC, 3SD. The numeral part indicates the number of gene states, and the alphabet indicates the mechanism by which NF-κB affect the gene transition rates. All reaction propensities are assumed to follow mass-action form.

c) **Initial conditions**

The single-cell measurements at time $t = 0$ after LPS stimulation in all conditions display some degree of expression for both genes. Thus, we model the joint distribution of gene state and mRNA copy number at the beginning of LPS stimulation as the solution of a time-invariant CME starting backward in time $-T_0$ where $T_0 \coloneqq 24$hr from the initial state with $G_0 = 1,\ G_1 = G_2 = \cdots = G_n = 0, R = 0$.

d) **Fitting single-gene models to observed marginal mRNA distributions under inhibitor-free condition**

We seek to match the model-predicted distributions of mRNA copy number to the measured distributions at five measurement times (0,0.5,1,2, and 4 hours) after LPS induction. Let $D_G^{\text{None}}$ denote the pooled dataset for inhibitor-free experiments that consist of single-cell observations of the form $(t, y)$ where $t$ is the time and $y$ the mRNA copy number of gene $G$ in the cell. We seek the parameter $\boldsymbol{\theta}_G$ that maximizes the log-likelihood of the form

$$\log L\left(D_G^{\text{None}} | \boldsymbol{\theta}_G^{\text{None}}\right) = \sum_{(t,y)\in D_G^{\text{None}}} \log p\left(t, y | \boldsymbol{\theta}_G^{\text{None}}\right).$$

e) **Optimization procedure for fitting single-gene dynamics to inhibitor-free data**

We use a parameter search strategy based implemented in the Python library PyGMO [4]. Global parameter search for each combination of model (2SA, 2SB, 3SA, 3SB, 3SC, 3SD) and gene (IL-1β, TNFα) followed the same routine that consists of two phases.

In the first phase, we conduct four global parameter searches based on the following heuristics: ant colony, differential evolution, particle swarm, and stochastic annealing with parallel independent chains. All four heuristics can be thought of as different ways to evolve a population of initial parameter guesses such that the next generation discovers better solutions than the previous one. We start each global search with 100 initial guesses sampled randomly from a uniform distribution supported by the bounding box of the parameters. Each population was evolved for up to 1000 generations, totaling about $4 \times 10^5$ loglikelihood evaluations per model and gene combination. After this phase, we pooled all the final populations of all heuristics and selected the best 100 parameter candidates for each gene-model combination.

In the second phase, we ran independent instances of the compass search algorithm starting from each of the parameter candidates selected from the first phase. After all of these searches concluded, we pooled all 100 optimal solutions found by all instances together and select the best parameter vector with the largest log-likelihood value.

f) **Three-state models provide superior fits to inhibitor-free expression data of both genes**

Supplementary Table 4 compares the models identified by fitting to the observed inhibitor-free marginal distributions of IL-1β and TNFα separately, using the optimization procedure described above. We use two criteria to compare how well the six models fit to the data. The first is the log-likelihood of the data given the model. The second is the Bayesian Information Criterion (BIC) defined for each model $\mathcal{M}$ as

$$\mathrm{BIC}(\mathcal{M}) = k_{\mathcal{M}} \log\left(\left|D_G^{\mathrm{None}}\right|\right) - 2\log\hat{L}_{\mathcal{M}},$$

where $k_{\mathcal{M}}$ is the number of parameters in the model (see Supplementary Table 1), $\left|D_G^{\mathrm{None}}\right|$ is the total number of single-cell observations used for fitting, and $\hat{L}_{\mathcal{M}}$ is the maximum value of the likelihood function given model $\mathcal{M}$.

For both genes, three variants of the three-state gene expression models, namely 3SA, 3SB, and 3SC, consistently rank as the top performers in terms of both performance metrics. For IL-1β, model 3SA (three-state, NF-κB enhances $G_0 \to G_1$ transition) performs best in terms of both metrics. The models 3SB and 3SC are the second and third best performers. For TNF-α, the best-performing model is 3SC (three-state, NF-κB enhances $G_1 \to G_2$ transition), but it is closely followed by 3SA and 3SB. Interestingly, the 3SD variant performs significantly worse than the rest of the proposed three-state models. In summary, three-state models perform significantly better than two-state models, even when their extra complexities have been taken into account.

### 3) Modeling joint observations of both genes across inhibitor conditions

a) **Combined mechanisms for jointly fitting and predicting mRNA distributions across genes and conditions**

We construct combined mechanistic models with the single-gene mechanisms described above as the basic building blocks. These models are assembled with the following principles. First, each gene is assigned one of the stochastic reaction network structures 2SA, 2SB, 3SA, 3SB, 3SC, 3SD (see Supplementary Table 2 and Supplementary Table 3 for specific reactions and parameters), so CMEs with the same network structure will describe the dynamics of each gene across conditions. We truncate the six possible mechanisms by half and select only the top performers (3SA, 3SB, 3SC) in terms of the total BIC computed from both IL-1β and TNFα observations under inhibitor-free condition (Supplementary Table

4). This results in nine possible combinations for consideration (Supplementary Figure 3). Second, we impose *parameter sharing* rules to reflect the effect of signals and inhibitors. In particular, the values of the parameters $r_1, r_2$ that describe NF-κB dynamics in eq.(1) are shared by the CMEs of all genes across all conditions. Across different drug inhibitor conditions, the CMEs for each gene share the same parameter vector except for the components that are modulated by NF-κB or inhibitors (see below).

b) **Modeling inhibitor effects**

We assume that different inhibitor treatments (MG132 or U0126) modulate the values of specific parameters in the models (Supplementary Table 5). Since U0126 inhibits C/EBP, which is assumed to increase the rate of the gene state transition from $G_0$ to $G_1$, we assume that the presence of U0126 alters the parameter $k_{01}$ in all six signal-modulated gene expression models. The inhibitor MG132 inhibits NF-κB, so the presence of this drug should alter the value of the signal strength parameter (see Supplementary Table 2 and Supplementary Table 3 for the specific parameter).

c) **Conditionally independent distributions product model**

The expression levels of the two genes across different conditions are assumed to be independent given the parameters describing NF-κB signal. Under this assumption, the distribution for the joint observation $\left(n_{IL-1\beta}^{\text{cond}}(t), n_{\text{TNF-}\alpha}^{\text{cond}}(t)\right)$ is modeled by a product of independent distributions as

$$\Pr\left(n_{\text{IL}-1\beta}^{\text{cond}}(t) = n_1, n_{\text{TNF}\alpha}^{\text{cond}}(t) = n_2 \middle| \boldsymbol{\theta}_{\mathcal{M}}\right) = p_{\mathcal{M}_1}\left(t, n_1 \middle| \boldsymbol{\theta}_{\text{IL}-1\beta}^{\text{cond}}\right) \cdot p_{\mathcal{M}_2}\left(t, n_2 \middle| \boldsymbol{\theta}_{\text{TNF}-\alpha}^{\text{cond}}\right) \qquad (2).$$

Here $\mathcal{M} = (\mathcal{M}_1, \mathcal{M}_2)$ could be one of the nine combinations

$$(A, A), (A, B), (A, C), (B, A), (B, B), (B, C), (C, A), (C, B), (C, C),$$

depending on which reaction network structures are chosen for IL-1β and TNF-α, respectively. The factor $p_{\mathcal{M}_1}\left(t, n_1 \middle| \boldsymbol{\theta}_{\text{IL}-1\beta}^{\text{cond}}\right)$ is the probability of observing $n_1$ IL-1β mRNA molecules at time $t$ under inhibitor condition $\text{cond} \in \{\text{None, MG132, U0126, Combined}\}$. It is obtained by solving the CME of the stochastic reaction network with structure $\mathcal{M}_1$ and parameter vector $\boldsymbol{\theta}_{\text{IL}-1\beta}^{\text{cond}}$. Similar interpretation applies to $p_{\mathcal{M}_2}\left(t, n_2 \middle| \boldsymbol{\theta}_{\text{TNF}\alpha}^{\text{cond}}\right)$.

The parameter vector $\boldsymbol{\theta}_{\mathcal{M}}$ collects all parameters necessary to derive the expression of all genes across all conditions. Its specific components depend on the model structures chosen for $\mathcal{M}$ and are summarized in Supplementary Table 6. Fix a model variant, the total parameter vector could be partitioned into

$$\boldsymbol{\theta}_{\mathcal{M}} = \left(\boldsymbol{\theta}_{\text{NFkB}}, \boldsymbol{\theta}_{\text{basal}}^{\text{IL}-1\beta}, b^{\text{IL}-1\beta,\text{None}}, b^{\text{IL}-1\beta,\text{MG}}, k_{01}^{\text{IL}-1\beta,\text{U0126}}, \boldsymbol{\theta}_{\text{basal}}^{\text{TNF}-\alpha}, b^{\text{TNF}-\alpha,\text{None}}, b^{\text{TNF}-\alpha,\text{MG}}, k_{01}^{\text{TNF}-\alpha,\text{U0126}}\right).$$

The reaction rates for the CME describing gene *S* under LPS induction without inhibitors are obtained from the parameter vector

$$\boldsymbol{\theta}_S^{\text{None}} = \left(r_1, r_2, \boldsymbol{\theta}_{\text{basal}}^T, b^S\right)^T.$$

On the other hand, the parameter vector to be fed into the CME for gene *S* under MG132 is then derived from the total parameter vector as

$$\boldsymbol{\theta}_S^{\text{MG}} = \left(r_1, r_2, \boldsymbol{\theta}_{\text{basal}}^T, b^{S,MG}\right)^T,$$

while the parameter vector $\boldsymbol{\theta}_S^{\text{U0126}}$ for the CME that describes gene S under U0126 is obtained by first copying the entries $r_1, r_2, \boldsymbol{\theta}_{\text{basal}}^T, b^S$ into their appropriate locations and replace the entry $k_{01}^S$ with $k_{01}^{S,\text{U0126}}$. The parameter vector $\boldsymbol{\theta}_S^{\text{MG+U0126}}$ for the CME that describes gene S under combined inhibitor treatment (MG132+U0126) is obtained by first copying $\boldsymbol{\theta}_S^{\text{MG}}$ and replace $k_{01}^S$ with $k_{01}^{S,\text{U0126}}$.

d) **Fitting joint observation models**

We seek parameters that maximize the likelihood of observing the mRNA copy numbers of both genes under inhibitor-free condition and single-inhibitor conditions (MG132 or U0126). Following eq.(2), the log-likelihood for observing the joint dataset for both genes at these three conditions are given by

$$\log L(D|\, \boldsymbol{\theta}_{\mathcal{M}})$$
$$= \sum_{\text{cond}} \sum_{(t,n_{\text{IL1}-\beta},n_{\text{TNF}-\alpha})\in D_{\text{cond}}} \log\left(p_{\mathcal{M}_1}\left(t, n_{\text{IL1}-\beta}\middle|\, \boldsymbol{\theta}^{\text{cond}}_{\text{IL1}-\beta}\right)\cdot p_{\mathcal{M}_2}\left(t, n_{\text{TNF}\alpha}\middle|\, \boldsymbol{\theta}^{\text{cond}}_{\text{TNF}-\alpha}\right)\right)$$
$$= \sum_{\text{cond}} \sum_{(t,n_{\text{IL1}-\beta},n_{\text{TNF}-\alpha})\in D_{\text{cond}}} \log\left(p_{\mathcal{M}_1}\left(t, n_{\text{IL1}-\beta}\middle|\, \boldsymbol{\theta}^{\text{cond}}_{\text{IL1}-\beta}\right)\right) + \log\left(p_{\mathcal{M}_2}\left(t, n_{\text{TNF}-\alpha}\middle|\, \boldsymbol{\theta}^{\text{cond}}_{\text{TNF}\alpha}\right)\right)$$

where the variable cond varies over the set of experimental conditions $\{None, MG132, U0126\}$.

e) **Optimization procedure for fitting to joint data across experiment conditions**

From the model selection results based on fits to the no-inhibitor data (see next section), we eliminate two-state mechanisms as well as the 3SD mechanism from consideration. Therefore, to model simultaneously fit the expression of both IL-1β and TNF-α in three conditions (No inhibitor, MG132, U0126) we consider only nine possible combinations, where we choose independently one of the mechanisms 3SA, 3SB, 3SC for each gene (IL-1β or TNF-α).

For each of the nine model structures, we run five independent stochastic annealing chains to attempt to find the global best fits. To make the starting guess for each chain, we copy the values of the best parameters found in the fit for inhibitor-free data to their corresponding slots in the combined model parameter vector. The initial guesses for the specific parameters for the two inhibitor experiments are simply copied from their counterparts in the inhibitor-free case.

f) **Model comparisons**

After obtaining the fits for all nine models, we evaluate their performance based on either how well they fit the “training” data (Supplementary Table 7, the “Fit Log-likelihood” column) and how well they predict the mRNA distributions under combined treatment (Supplementary Table 7, the “Prediction Log-likelihood” column). Three models (3SCA, 3SBA, and 3SAA) stand out in terms of these two metrics (Supplementary Figure 4). Of these three, the 3SCA variant has the highest total log-likelihood (the sum of fit log-likelihood and prediction log-likelihood). To rule out the possibility that one of the previously 27 discarded combinatorial models could have outperformed the final models, we compared the ability to predict IL-1β and TNFα distributions in the inhibitor-free dataset of the nine combined models against the six independent gene expression models (Supplementary Figure 6). The two models 3SCA and 3SBA still outperform the single-gene mechanisms we previously discarded despite having been constrained by sharing the parameters between genes and conditions and having to fit to more data from the inhibitory treatment experiments.

**Supplementary Tables**

| Model | NF-κB affects | Number of gene states | Number of parameters |
|---|---|---|---|
| 2SA | $G_0 \to G_1$ (increase) | 2 | 8 |
| 2SB | $G_1 \to G_0$ (decrease) | 2 | 8 |
| 3SA | $G_0 \to G_1$ (increase) | 3 | 11 |
| 3SB | $G_1 \to G_0$ (decrease) | 3 | 11 |
| 3SC | $G_1 \to G_2$ (increase) | 3 | 11 |
| 3SD | $G_2 \to G_1$ (decrease) | 3 | 11 |

***Supplementary Table 1.*** *Number of parameters in the six proposed mechanisms for single-gene transcription dynamics. The models differ in the number of gene states and in how NF-κB signal affects gene transitions.*

| Reaction | Basal | 2SA | 2SB |
|---|---|---|---|
| $G_0 \to G_1$ | $k_{01}$ | $k_{01} + b_{01} S_{\mathrm{NF-\kappa B}}(t)$ | X |
| $G_1 \to G_0$ | $k_{10}$ | X | $\max(0, k_{10} - b_{10} S_{\mathrm{NF-\kappa B}}(\mathrm{t}))$ |
| $G_0 \to \mathrm{RNA}$ | $\alpha_0$ | X | X |
| $G_1 \to \mathrm{RNA}$ | $\alpha_1$ | X | X |
| $\mathrm{RNA} \to 0$ | $\delta$ | X | X |

***Supplementary Table 2.*** *Reactions and reaction rates for the four NF-κB-activated two-state gene expression reaction networks considered in this paper. The notation "X" means that the parameter is shared with the basal model. The two variants 2Sx (x is either A or B) differ in the reaction rate that is modulated by NF-κB. The multiplicative constants $b_{ij}$ represent the level of influence the signal has on the transition rate from gene state i to gene state j.*

| Reaction | Basal | 3SA | 3SB | 3SC | 3SD |
|---|---|---|---|---|---|
| $G_0 \rightarrow G_1$ | $k_{01}$ | $k_{01} + b_{01} S_{\mathrm{NF-\kappa B}}(t)$ | X | X | X |
| $G_1 \rightarrow G_0$ | $k_{10}$ | X | $\max(0, k_{10} - b_{10} S_{\mathrm{NF-\kappa B}}(\mathrm{t}))$ | X | X |
| $G_1 \rightarrow G_2$ | $k_{12}$ | X | X | $k_{12} + b_{12} S_{\mathrm{NF-\kappa B}}(t)$ | X |
| $G_2 \rightarrow G_1$ | $k_{21}$ | X | X | X | $\max(0, k_{21} - b_{21} S_{\mathrm{NF-\kappa B}}(\mathrm{t}))$ |
| $G_0 \rightarrow$ RNA | $\alpha_0$ | X | X | X | X |
| $G_1 \rightarrow$ RNA | $\alpha_1$ | X | X | X | X |
| $G_2 \rightarrow$ RNA | $\alpha_2$ | X | X | X | X |
| RNA $\rightarrow 0$ | $\delta$ | X | X | X | X |

***Supplementary Table 3.*** *Reactions and reaction rates for four NF-κB-activated three-state gene expression reaction networks considered in this paper. The notation "X" means that the parameter is shared with the basal model. The four variants 3Sx (x is either A, B, C, or D) differ in the reaction rate that is modulated by NF-κB. The multiplicative constants $b_{ij}$ represent the level of influence the signal has on the transition rate from gene state i to gene state j. In particular, the larger $b_{ij}$ is, the larger the change in gene state transition rate in response to NF-κB concentration.*

| Model | IL-1β | | TNF-α | |
|---|---|---|---|---|
| | log-likelihood | BIC | log-likelihood | BIC |
| 2SA | -71606.16 | 143287.22 | -61234.15 | 122543.19 |
| 2SB | -72145.63 | 144366.14 | -63172.48 | 126419.86 |
| 3SA | **-71374.57** | **142852.12** | -60818.20 | 121739.37 |
| 3SB | -71441.88 | 142986.74 | -61086.19 | 122275.35 |
| 3SC | -71453.01 | 143008.993 | **-60808.89** | **121720.75** |
| 3SD | -72314.90 | 144732.77 | -63271.87 | 126646.71 |

***Supplementary Table 4.*** *Comparison of different proposed mechanisms in terms of log-likelihood and Bayesian Information Criterion (BIC) using data collected from inhibitor-free experiments. The best performing model for each column is highlighted in boldface.*

| Mechanism / Inhibitor | 2SA or 3SA | 2SB or 3SB | 3SC | 3SD |
|---|---|---|---|---|
| MG132 | $b_{01}$ | $b_{10}$ | $b_{21}$ | $b_{12}$ |
| U0126 | $k_{01}$ | | | |

***Supplementary Table 5.*** *Description of which parameters are altered by inhibitor treatment in each model. The effect of small-molecule inhibitors MG132 and U0126 are assumed to change the value of specific parameters in the reaction network. Because signals modulate the reaction rates of the basal gene-expression model differently in different signal-activated models, the inhibitors also alter different parameters depending on the model considered.*

| | A | B | C | Interpretation |
|---|---|---|---|---|
| 1 | $r_1$ | X | X | NF-κB signal |
| 2 | $r_2$ | X | X | |
| 3 | $k_{01}^{\mathrm{IL-1\beta}}$ | X | X | Basal IL-1β expression |
| 4 | $k_{10}^{\mathrm{IL-1\beta}}$ | X | X | |
| 5 | $k_{12}^{\mathrm{IL-1\beta}}$ | X | X | |
| 6 | $k_{21}^{\mathrm{IL-1\beta}}$ | X | X | |
| 7 | $\alpha_0^{\mathrm{IL-1\beta}}$ | X | X | |
| 8 | $\alpha_1^{\mathrm{IL-1\beta}}$ | X | X | |
| 9 | $\alpha_2^{\mathrm{IL-1\beta}}$ | X | X | |
| 10 | $\delta^{\mathrm{IL-1\beta}}$ | X | X | |
| 11 | $b_{01}^{\mathrm{IL-1\beta,None}}$ | $b_{10}^{\mathrm{IL-1\beta,None}}$ | $b_{12}^{\mathrm{IL-1\beta,None}}$ | NF-κB modulated IL-1β gene state switching rate (without inhibitor) |
| 12 | $b_{01}^{\mathrm{IL-1\beta},\,MG}$ | $b_{10}^{\mathrm{IL-1\beta},\,MG}$ | $b_{12}^{\mathrm{IL-1\beta},\,MG}$ | NF-κB modulated IL-1β gene state switching rate (with MG132) |
| 13 | $k_{01}^{\mathrm{IL-1\beta,\,U0126}}$ | X | X | U0126-modulated gene activation rate for IL-1β |
| 14 | $k_{01}^{\mathrm{TNF\alpha}}$ | X | X | basal TNF-α expression |
| 15 | $k_{10}^{\mathrm{TNF\alpha}}$ | X | X | |
| 16 | $k_{12}^{\mathrm{TNF\alpha}}$ | X | X | |
| 17 | $k_{21}^{\mathrm{TNF\alpha}}$ | X | X | |
| 18 | $\alpha_0^{\mathrm{TNF\alpha}}$ | X | X | |
| 19 | $\alpha_1^{\mathrm{TNF\alpha}}$ | X | X | |
| 20 | $\alpha_2^{\mathrm{TNF\alpha}}$ | X | X | |
| 21 | $\delta^{\mathrm{TNF\alpha}}$ | X | X | |
| 22 | $b_{01}^{\mathrm{TNF-\alpha,None}}$ | $b_{10}^{\mathrm{TNF-\alpha,None}}$ | $b_{12}^{\mathrm{TNF-\alpha,None}}$ | NF-κB modulated TNF-α gene state switching rate (without inhibitor) |
| 23 | $b_{01}^{\mathrm{TNF-\alpha},\,MG}$ | $b_{10}^{\mathrm{TNF-\alpha},\,MG}$ | $b_{12}^{\mathrm{TNF-\alpha},\,MG}$ | NF-κB modulated TNF-α gene state switching rate (with MG132) |
| 24 | $k_{01}^{\mathrm{TNF-\alpha,\,U0126}}$ | X | X | U0126-modulated gene activation rate for TNF-α |

***Supplementary Table 6.*** *Interpretation of parameters for the combined gene state model variants. There are 9 possible model variants, depending on the reaction network structures chosen independently for IL-1β and TNF-α. Entries marked with "X" have the same values with its counterpart in column "A". To obtain the parameter vector for a combination, concatenate the parameters on the appropriate columns. For example, for the parameter vector of "BA", concatenate the parameters 1-2 on the first column, parameters 3-13 on column "B", and parameters 14-24 on column "A".*

| Model | Fit Log-likelihood | Prediction Log-likelihood | Total Log-likelihood (both fit and prediction) |
|---|---|---|---|
| 3SAA | -354648.90 | -85201.88 | -439850.77 |
| 3SAB | -355948.23 | -85923.22 | -441871.46 |
| 3SAC | -356952.18 | -86782.73 | -443734.90 |
| 3SBA | **-354255.62** | -85238.98 | -439494.60 |
| 3SBB | -355727.55 | -85956.85 | -441684.40 |
| 3SBC | -355605.22 | -86235.61 | -441840.83 |
| 3SCA | -354338.45 | **-85069.20** | **-439407.65** |
| 3SCB | -355754.57 | -85949.04 | -441703.62 |
| 3SCC | -356341.18 | -86528.21 | -442869.39 |

***Supplementary Table 7.*** *Comparison between different models for fitting IL-1β and TNF-α in the drug-free and single-inhibitor conditions (second column), prediction of the joint IL-1β and TNF-α response in the combined inhibitor condition (third column), and sum of the fit and prediction loglikelihoods (right-most column) .*

### Supplementary Figures

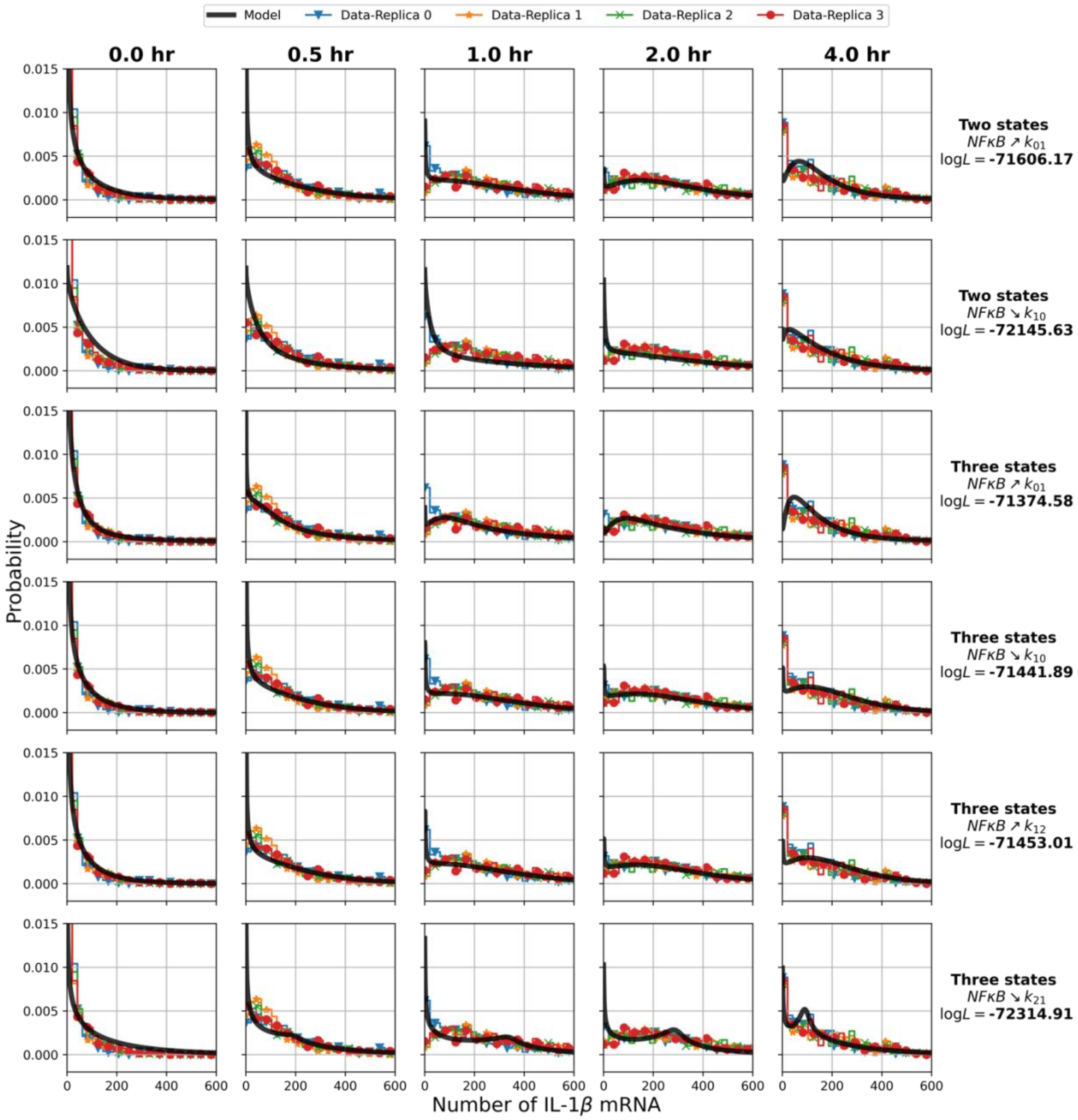

***Supplementary Figure 1***.*Fits for IL-1β expression under no inhibitor treatments.*

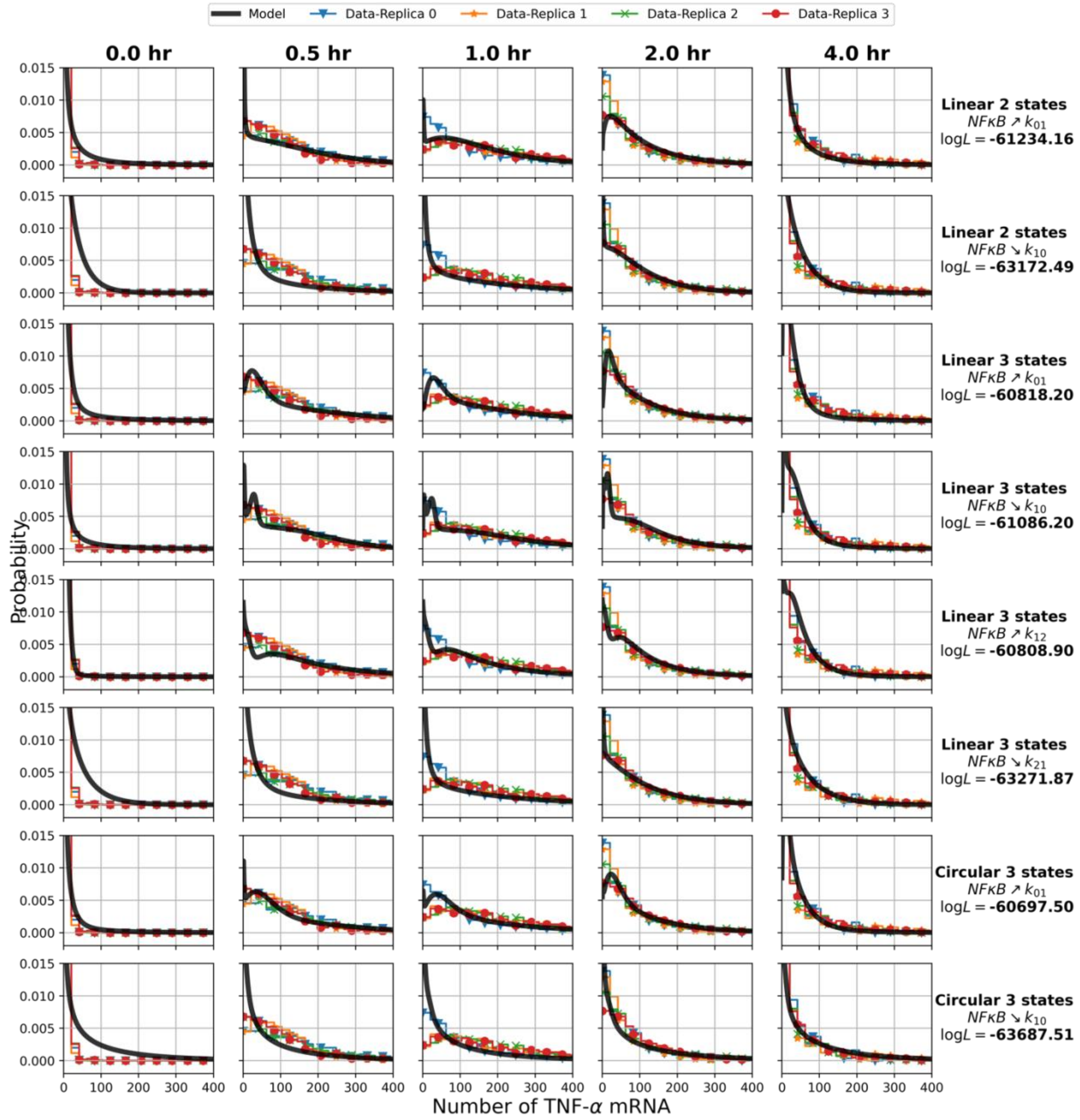

***Supplementary Figure 2.*** *Fits for TNF-$\alpha$ expression under no inhibitor treatments.*

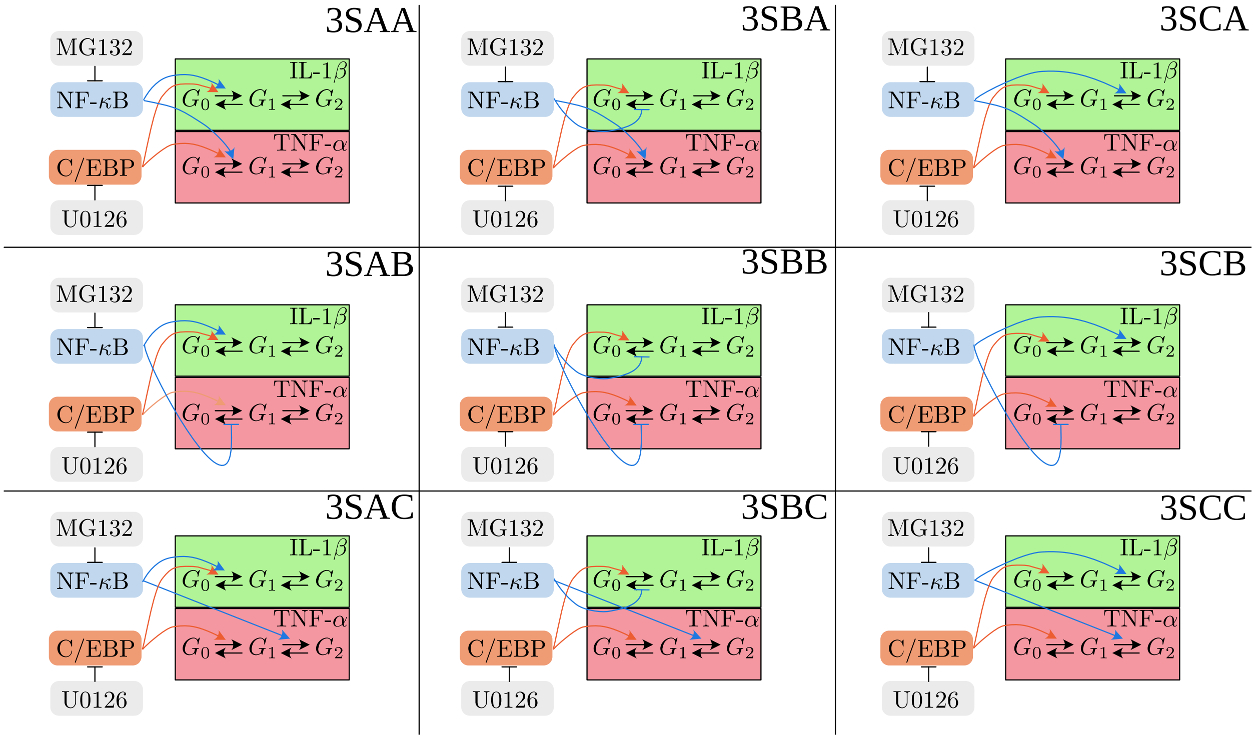

***Supplementary Figure 3.*** *Combined three-state models for fitting the expression of IL-1β and* TNF$\alpha$ *expression.*

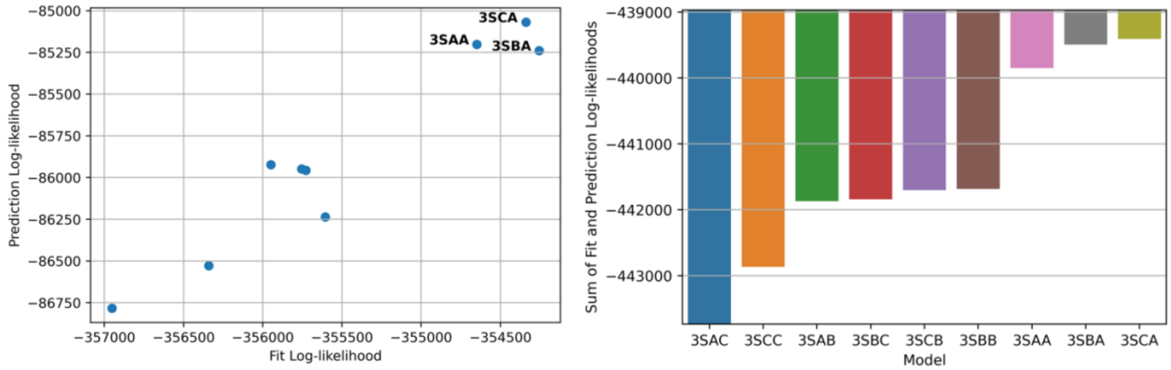

***Supplementary Figure 4.*** *Performance evaluation for simultaneous two-gene, multi-condition models based on their abilities to explain mRNA distributions of both genes under inhibitor-free, MG132 and U0126 (which were used for fitting the parameters), and to predict mRNA distributions under combined MG132+U0126 treatment (which were not used for parameter fitting).*

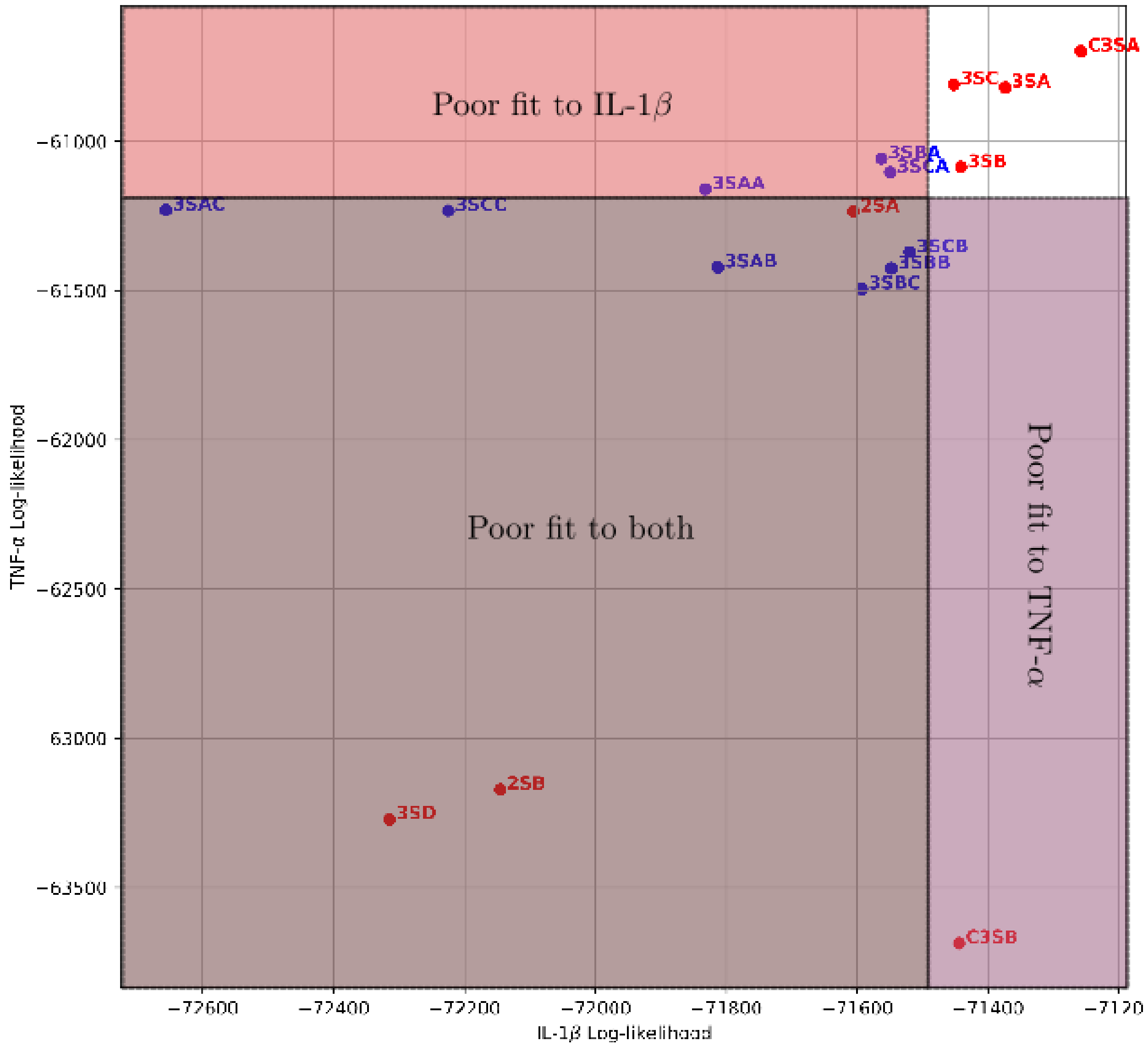

***Supplementary Figure 5.*** *Performance of models for fitting to expression data of IL-1β and TNFα under LPS stimulation without inhibitor treatments. The horizontal and vertical axes represent log-likelihoods of observed IL-1β and TNFα data given the models. The red dots represent the independent expression models that are fit independently to the observed marginal distributions of both genes separately. The blue dots represent combined models with parameter sharing constraints and were fit to IL-1β and TNFα under three experimental conditions (No inhibitors, MG132, U0126) simultaneously.*

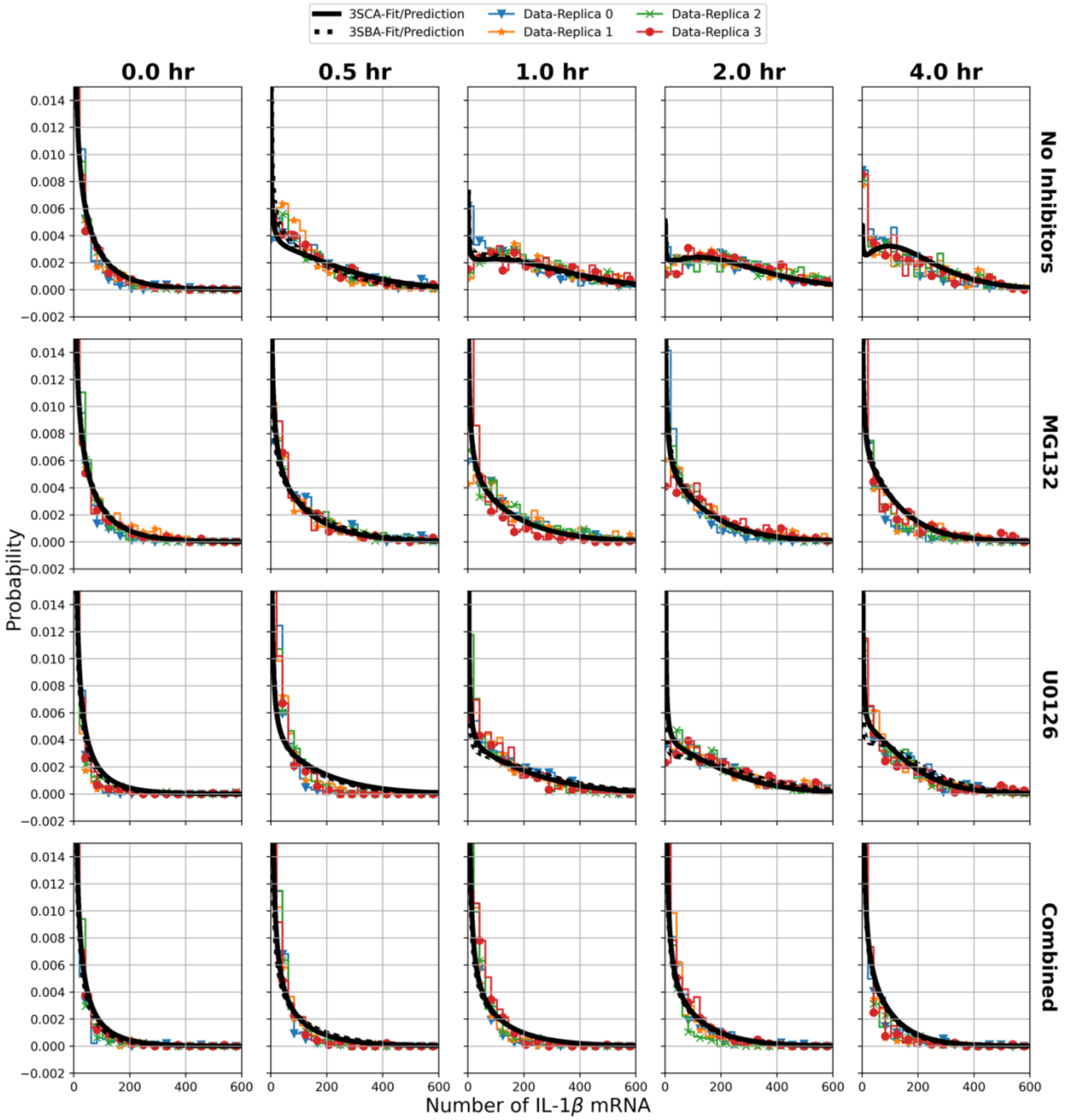

**Supplementary Figure 6.** *Fits and predictions made by the single-gene model for the distributions of IL-1β mRNA copy number at four different experimental conditions (top to bottom: No inhibitors, MG132, U0126, combined MG132+U0126) and five measurement time points (left to right: 0 hr, 30 min, 1 hr, 2 hr, 4 hr). The black solid lines represent the predictions made by the single-gene model (Fig 3 in the main text) using parameters fitted to measured IL-1β mRNA distributions at three conditions: No Inhibitors, MG132, U0126. These parameters are then combined to predict the transcriptional response under the application of both inhibitors (bottom row). Measured distributions from four independent biological replicas are represented as marked lines.*

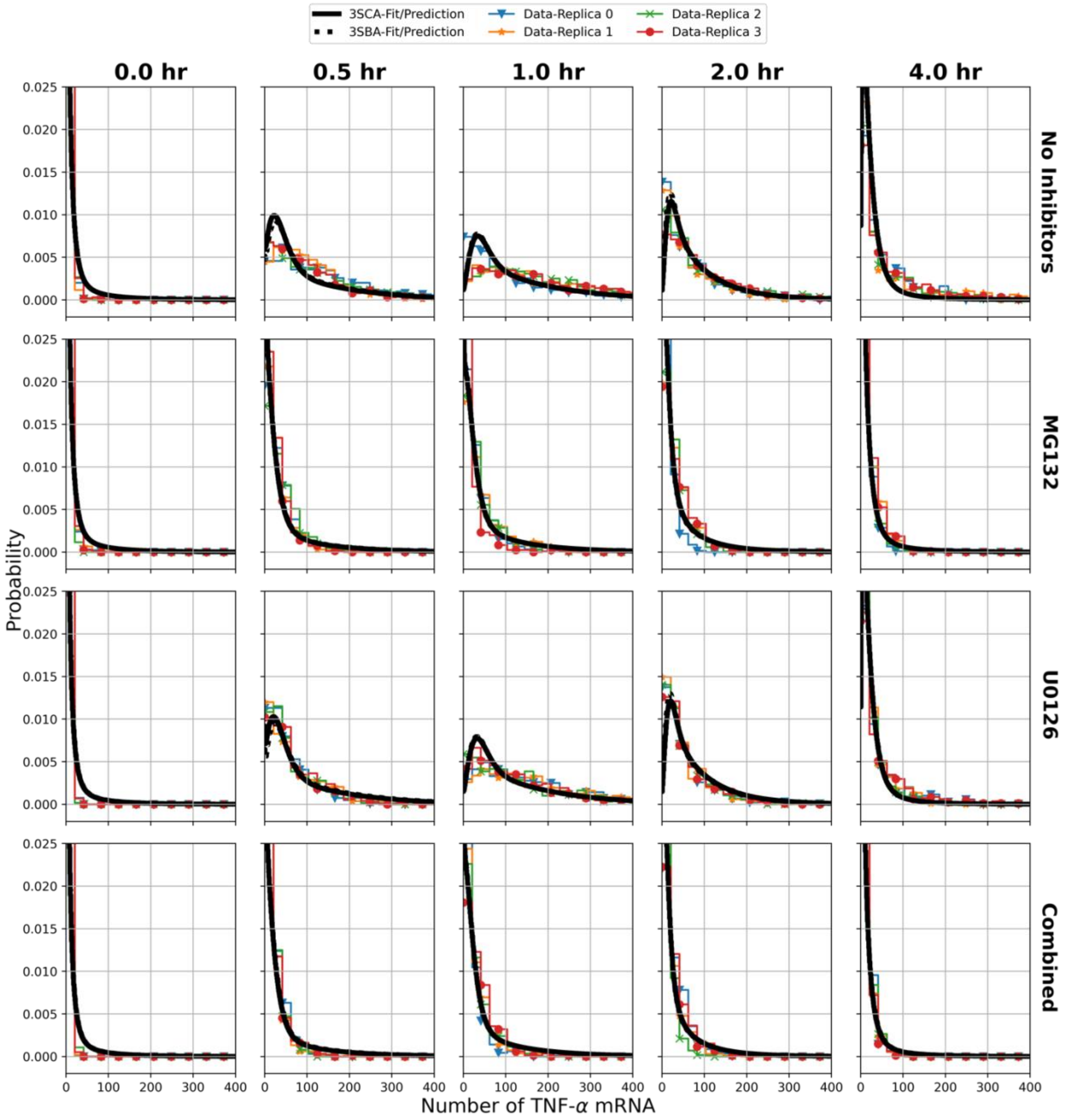

**Supplementary Figure 7.** *Fits and predictions made by the single-gene model for the distributions of TNF-α mRNA copy number at four different experimental conditions (top to bottom: No inhibitors, MG132, U0126, combined MG132+U0126) and five measurement time points (left to right: 0 hr, 30 min, 1 hr, 2 hr, 4 hr). The black solid line represents the prediction made by the single gene model (Fig 3 in the main text) using parameters fitted to measured TNF-α mRNA distributions at three conditions: No Inhibitors, MG132, U0126. These parameters are then combined to predict the transcriptional response under the application of both inhibitors (bottom row). Measured distributions from four independent biological replicas are represented as marked lines.*
